# A transcriptional and regulatory map of mouse somitogenesis

**DOI:** 10.1101/2023.01.24.525253

**Authors:** Ximena Ibarra-Soria, Elodie Thierion, Gi Fay Mok, Andrea E. Münsterberg, Duncan T. Odom, John C. Marioni

## Abstract

The mammalian body plan is shaped by rhythmic segmentation of mesoderm into somites, which are transient embryonic structures consisting of hundreds of cells that form down each side of the neural tube. We have systematically analysed the genome-wide transcriptional and chromatin dynamics occurring within nascent somites, from early inception of somitogenesis to the latest stages of body plan establishment. We created matched gene expression and open chromatin maps for the three leading pairs of somites at six time points during embryonic development. Here we show that the rate of somite differentiation accelerates as development progresses. We identified a conserved maturation programme followed by all somites after segmentation, but somites from more developed embryos concomitantly switch on differentiation programmes from derivative cell lineages soon after segmentation. Integrated analysis of the somitic transcriptional and chromatin activities revealed opposing regulatory modules controlling the onset of differentiation. We identified transcription factors expressed during early development that inhibit the activity of proteins required for commitment and differentiation of skeletal cell populations. Our results provide a powerful, high-resolution view of the molecular genetics underlying somitic development in mammals.

## INTRODUCTION

The segmentation of the body plan during early embryogenesis is a fundamental and conserved feature of all vertebrate species. It results in the metameric organisation of the vertebrae and the associated skeletal muscles, nerves, and blood vessels. This segmentation is established via formation of somites, which are transient embryonic structures consisting of hundreds of cells that bud off from the anterior tip of the presomitic mesoderm (PSM) on each side of the neural tube. Each pair of somites is symmetrically and rhythmically formed along the anterior-posterior axis according to the *clock and wavefront model (Cooke and Zeeman 1976)*.

This model integrates spatiotemporal information from waves of transcriptionally oscillating genes in the PSM (the molecular clock) and antagonistic signalling gradients along the embryo axis (the wavefront). The molecular oscillator is known as the segmentation clock, which drives cyclic and synchronised gene expression along the PSM (Palmeirim et al. 1997). The so-called *clock genes* belong to the Notch, Wnt and FGF signalling pathways (Dequéant and Pourquié 2008). The wavefront involves posterior gradients of Wnt and FGF signalling that are counteracted by an opposing gradient of retinoic acid secreted from the somites (Bénazéraf and Pourquié 2013). When the segmentation clock reaches cells that have passed the wavefront, segmentation genes, including *Mesp2*, are activated, leading to the specification of the somite boundary (Saga 2012). As well as specifying the somite boundaries, retinoic acid signalling suppresses signals that break left-right symmetry, ensuring that somite production is bilaterally symmetric (J. Vermot and Pourquié 2005; Julien Vermot et al. 2005). This periodic addition of somites underlies body plan generation in all vertebrates, and the oscillating signals from Notch, Wnt and FGF pathways are conserved in the PSM of model organisms as diverse as mouse, chicken, and zebrafish (Krol et al. 2011).

The specification of somites along the anterior-posterior axis is determined before somitogenesis by Hox gene expression (Krumlauf 1994) and the specific combination of Hox genes expressed along the axis establishes the identity of the resulting vertebrae (Wellik 2007). Somites are further patterned along the dorso-ventral and medio-lateral axes, giving rise to two somitic derivatives found in all vertebrates: the sclerotome (precursor of vertebral and rib cartilage, tendons, and blood vessels) and the dermomyotome (precursor of skeletal muscles and back dermis). Fate specification to either derivative is controlled by signals from adjacent tissues. Ventral cells of the somite differentiate into sclerotome under the influence of Shh signals from the notochord and the floor-plate of the neural tube. Dorsal cells instead receive Wnt signals from the neural tube and the ectoderm and BMP4 from the lateral mesoderm, to give rise to the dermomyotome (Weldon and Münsterberg 2022).

Master transcriptional regulators driving somite differentiation have been identified through classical genetic approaches (Yusuf and Brand-Saberi 2006; Christ, Huang, and Scaal 2007). However, how these master regulators orchestrate somitogenesis through embryonic space and time, and indeed what genes they directly regulate, remains less clear. While several studies have used gene expression microarrays to characterise gene expression patterns during somitogenesis, these studies have all been performed in the presomitic mesoderm (Krol et al. 2011; Dequéant et al. 2006; Ozbudak, Tassy, and Pourquié 2010).

Here, we map the transcriptional and chromatin changes that occur across somitogenesis by performing high-resolution RNA and ATAC-sequencing of individual, manually microdissected somites at six developmental stages. By comparing the three most recently segmented somites, we characterised the molecular basis of the earliest stages of somite maturation. Additionally, we identified patterns of dynamic regulatory activity across development, with pronounced differences between somites that give rise to differing types of vertebrae. By characterising the biological processes dominating each stage, we found that somite differentiation accelerates with developmental progression. Finally, we used the combined information from the transcriptional and chromatin maps to define regulatory modules with differing activity during early and late development. These molecular programmes control the onset of differentiation, thus regulating the timing of skeletal system development.

## RESULTS

### A high resolution transcriptional and regulatory map of somitogenesis

To characterise the transcriptional changes that orchestrate mouse somitogenesis, we generated coupled transcriptional and chromatin accessibility profiles of individual somite pairs, across embryonic development. Each somite typically contains 500-1000 cells, which is sufficient to generate high-resolution small bulk data. We first compared the transcriptomes of matched left and right somites dissected from 20-25 somite embryos, and observed no significant differences in expression (Figure S1), indicating that from a molecular genetics perspective, the two somites were indistinguishable. Therefore, for each somite pair, we used one somite to map the transcriptome (RNA-seq) and one to map matched open chromatin (ATAC-seq) (Figure 1A).

**Figure 1.**
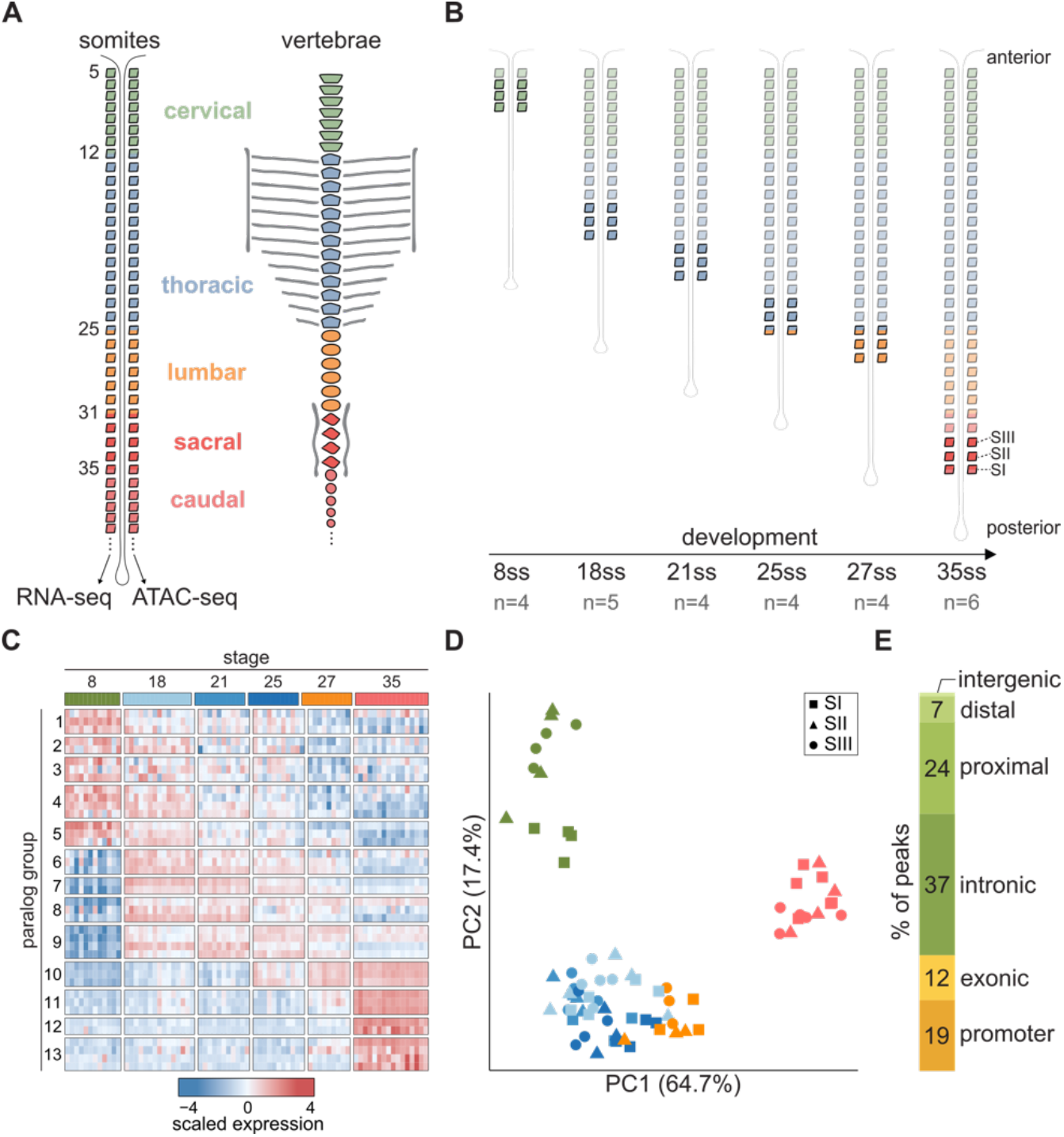
Expression and chromatin profiling of mouse somitogenesis. **A)** Schematic of somite pairs on each side of the neural tube, and the corresponding vertebrae structures they will form. One somite from each pair was used for RNA-seq, and the other for ATAC-seq. Somites and vertebrae are coloured based on their vertebral identity (cervical, thoracic, lumbar, sacral or caudal). Somites at boundaries between two different types of vertebrae are numbered. The first four somite pairs (occipital) are not shown. **B)** Somites collected in this study. From each embryo, the three most posterior somites (SI-SIII) were collected. Embryos profiled were from six different developmental stages, determined by the number of somites. ss = somite stage. n = number of embryos collected. **C)** Heatmap of the expression of all Hox genes. Samples are ordered in columns according to their observed somite stage. Each row is a different Hox gene, ordered by paralogous groups from 1 to 13. Expression is represented as z-scores. **D)** Principal component analysis of the expression of Hox genes orders somites consistently with their observed somite stage. **E)** Proportion of open chromatin regions classified based on their genomic context.

After segmentation, somites maintain a round shape for several hours before undergoing an epithelial-to-mesenchymal transition (EMT) when cells commit to somite-derived lineages and initiate migration (Yusuf and Brand-Saberi 2006). To study the molecular changes associated with fate commitment, we collected the three most posterior pairs of somites, which correspond to those most recently segmented, and that have not yet begun EMT (Christ, Huang, and Scaal 2007; M. Jacob, Christ, and Jacob 1975) (Figure 1B). To understand how somitogenesis progresses across embryonic development, we sampled these somite trios from embryos at six different developmental stages. We defined the embryonic stage by counting the total number of somite pairs, and profiled at least four different embryos containing 8, 18, 21, 25, 27 and 35 pairs of somites (Figure 1B and Table S1). These stages span four of the five different types of vertebrae (cervical, thoracic, lumbar and sacral; Figure 1A), providing profiles of somites that will contribute to all four structures.

We generated matched transcriptome (RNA-seq) and open chromatin (ATAC-seq) maps from the vast majority (71/81) of the samples (Tables S1). From the 77 RNA-seq libraries, all but one produced good-quality transcriptomes (Tables S2). We normalised for sequencing depth and corrected for batch effects associated with the date of somite collection (Methods; Figure S2).

Somites from specific axial levels express particular combinations of Hox genes (Wellik 2007), which are directly associated with segment identity (Kmita and Duboule 2003; Mallo, Wellik, and Deschamps 2010). Somites from different embryonic stages consistently showed clear differences in the class and expression level of Hox genes (Figure 1C), and the expression of Hox genes alone accurately ordered samples according to our observed somite stage (Figure 1D).

ATAC-seq libraries were successfully produced from 75 samples, but 25 of these were removed after applying stringent quality control criteria (Figure S3 and Table S3). The remaining 50 open chromatin maps showed efficiency biases, which were correlated with mean fragment abundance (Figure S4A-B). To compensate for this trend, we used a loess-based normalisation strategy (Figure S4C). Additionally, we applied the same batch correction approach that was used on the RNA-seq data to remove technical variation (Figure S4D-E).

We classified the possible functional role of open chromatin regions based on their genomic location. Peaks that were within 200 bp of an annotated transcription start site were deemed promoter-like elements and represent 19% of total peaks; an additional 12% of peaks overlapped gene exons. The remainder of the peaks were annotated as enhancer-like elements, and subdivided into *proximal* (24%) or *distal* (7%) if they were within 25 and 100 kb of an annotated gene, respectively; or *intergenic* (1%) (Figure 1E).

### Epithelial somites deploy a shared maturation programme across embryonic development

We systematically profiled the three most recently segmented somites, which are at the beginning of the differentiation process that will give rise to all somitic derivatives, including muscle, bone, cartilage and dermis (H. J. Jacob, Christ, and Jacob 1974; Christ et al. 1992; Ordahl and Le Douarin 1992; Aoyama 1993). Following the nomenclature proposed by Christ and Ordahl (Christ and Ordahl 1995), we refer to each somite in these trios as somites I, II and III, from the most posterior to the most anterior, respectively (Figure 2A).

**Figure 2.**
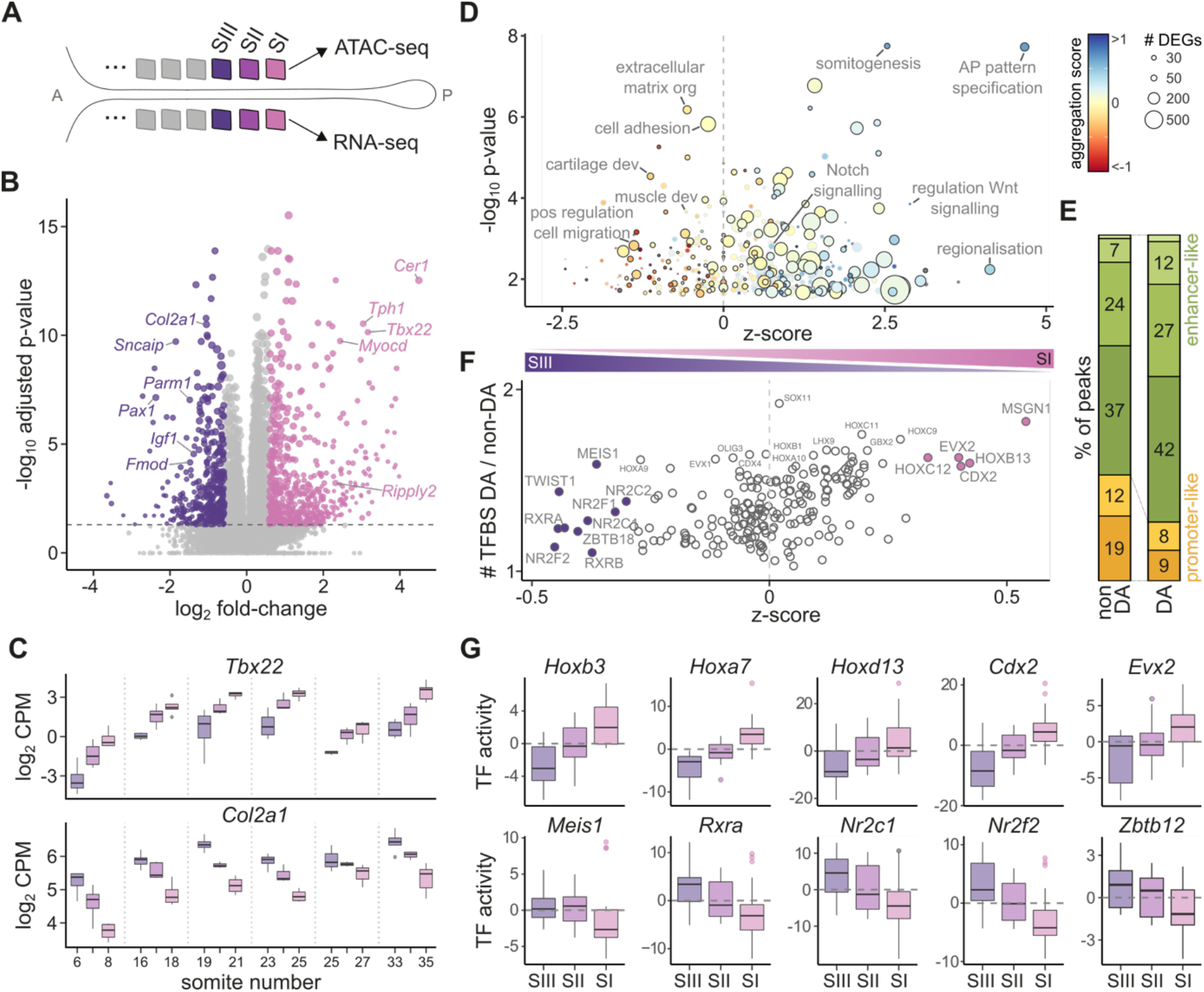
Somites follow a conserved maturation programme across development. **A)** Schematic indicating the somite trios profiled from each embryo. A: anterior; P: posterior. Colour and shading scheme is preserved throughout all figures to indicate the different somites. **B)** Volcano plot of expression changes between somites I and III. Genes significantly differentially expressed are coloured. **C)** Differences in gene expression for two representative genes (*Tbx22* and *Col2a1*) among somites I,II,III are consistently maintained across developmental stages. **D)** Significantly enriched Gene Ontology functional categories in the set of differentially expressed genes. Enrichment significance is shown on the y axis. The x-axis indicates whether a term contains a majority of genes that are downregulated (positive) or upregulated (negative). Points are coloured based on an ‘aggregation score’, which corresponds to the average fold-change of all differentially expressed genes in the GO term. The size of the points indicates the number of differentially expressed genes in each term. Outlined points correspond to terms that are also significantly enriched in the set of differentially accessible chromatin regions. **E)** Barplot of the proportion of peaks falling in different genomic contexts. Differentially accessible (DA) regions between somites I,II,III are enriched for enhancers. Colours indicate the same classes as in Figure 1E. **F)** Similar to D but showing the enrichment of transcription factor binding sites (TFBS) in differentially accessible peaks. **G)** Representative examples of motif activity dynamics for TFs that are enriched in differentially accessible peaks. Positive (negative) activity scores indicate the regions are more (less) accessible compared to background chromatin. Hox and other hoemeodomain TF binding sites close in mature somites, while C4 zinc finger class of receptors (*Rxra, Nr2c1, Nr2f2, Zbtb12*) sites become more accessible in SIII.

To characterise the molecular changes underlying somite maturation, we compared all pairwise combinations of somites I, II and III at each developmental stage. We identified a median of 453 genes that significantly differ per stage (FDR < 5% and |fold-change|>1.5). Most differentially expressed genes had subtle changes in expression, with half showing less than a two-fold difference between any two somites. To increase statistical power, we repeated the analysis using samples from different stages as replicates, and detected genes that showed consistent changes regardless of developmental stage. Altogether, we identified 2,977 significantly differentially expressed genes. Similar numbers of genes were up and down-regulated, with the strongest differences manifested between somites I and III (Figure 2B). The vast majority of differentially expressed genes (75.8%) showed consistent expression dynamics across different stages. However, most genes (86.9%) also showed differences in expression levels across developmental time, illustrating the complex regulatory dynamics prevalent during embryonic development (Figure 2C).

The genes downregulated along somite maturation were enriched for biological processes related to regionalisation and pattern specification, which are active in the PSM and lead to somite segmentation. Consistently, both the Wnt and Notch signalling pathways were preferentially downregulated (Dequéant and Pourquié 2008) (Figure 2D). In contrast, a steady progression towards EMT during somitogenesis was reflected by the upregulation of cell adhesion and migration programs, together with a switch to positive regulation of Rho and ERK signalling (Figure 2D).

The open chromatin landscape was similarly dynamic across the somite trios, with 2,701 genomic regions showing significantly different accessibility levels (FDR < 5% and |fold-change|>1.5). Open chromatin regions that actively changed between somites were enriched for enhancer-like regions, with fewer promoter elements (Figure 2E). Indeed, only 506 (18.7%) differentially accessible regions were located within 5 kb of a differentially expressed gene, indicating that the regulatory mechanisms driving expression changes operate through distal regulatory elements, rather than by directly modulating chromatin accessibility at promoters. The coordination of dynamic chromatin accessibility and gene expression changes was also reflected in their shared over-representation of the same biological functions (Figure 2D).

Next, we annotated transcription factor (TF) binding motifs within ATAC-seq peaks and identified 201 regulators whose binding motifs were significantly enriched in the dynamic regions, when compared to static open chromatin (Figure 2F). These included Hox factors, as well as multiple members of the homeodomain, Tal, Sox and NK families. For example, binding motifs for MSGN1 were present in 22% of all dynamic regions (compared to 12% in non differentially accessible chromatin), and most of these peaks showed reduced accessibility in more mature somites, consistent with the role of this protein as a master regulator of PSM differentiation (Chalamalasetty et al. 2014) (Figure 2F). In contrast, the dynamic peaks with binding motifs for TWIST1, a critical factor mediating EMT, were more accessible in the most mature somites (Lamouille, Xu, and Derynck 2014) (Figure 2F).

Finally, to understand how the overrepresented transcription factors regulate somite maturation, we analysed the accessibility dynamics of the genomic loci with binding sites for each of these TFs. Binding sites for all Hox proteins, regardless of their paralogous group or stage activity pattern, showed decreased accessibility upon somite maturation (Figure 2G). This behaviour also extended to most of the other TFs enriched in differentially accessible peaks (Figure 2G). One notable exception gained accessibility as somites matured: the C4 zinc finger class of receptors, which includes the retinoid X receptor-related factors, critical in mediating the biological effects of retinoid signalling and its differentiation-inducing activity (Draut, Liebenstein, and Begemann 2019) (Figure 2G).

### Molecular remodelling across development regulates somite responses to the signalling environment

Our data revealed profound changes in transcriptional and regulatory activity in somites I-III across development. We compared the RNA-seq profiles of somites among all different developmental stages (Figure 3A) and identified 10,691 genes with significant changes in expression (FDR < 5% and |fold-change|>1.5; see Methods for details; Figure 3B), including most known transcription factors (838 from a total of 1,310 expressed). The chromatin landscape was also remodelled extensively, with 33,013 open chromatin regions showing significant differences in accessibility (Figure S5A). In contrast to the changes observed across somite maturation, a much higher proportion of differentially accessible loci were in promoters or close to differentially expressed genes (Figure S5B), indicating that across development widespread chromatin remodelling plays a crucial role in controlling the genes available for expression.

**Figure 3.**
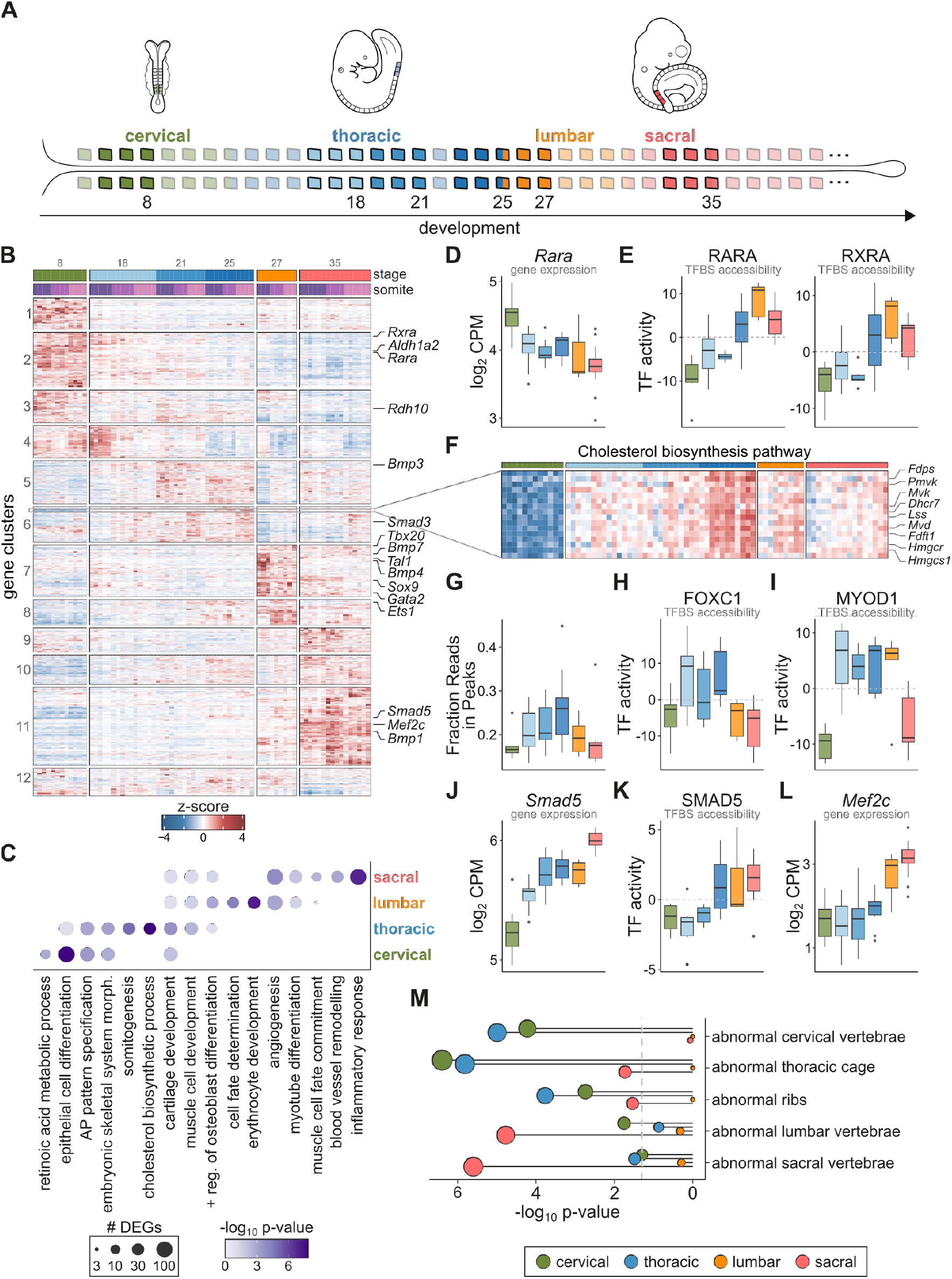
Epithelial somites at late development activate differentiation programmes of derivative lineages absent in early stages. **A)** Schematic indicating the somites profiled across development, and their vertebral fate. Colour scheme is preserved throughout all figures to indicate the different stages. **B)** Heatmap of expression of genes differentially expressed across development. Samples (columns) are ordered based on their somite number, and their stage and somite level are indicated at the top. **C)** Gene ontology term enrichment analysis results for sets of genes with highest activity at particular vertebral fates. The size of the circles indicates the number of differentially expressed genes in each term-fate combination, and the intensity of the colour corresponds to the significance of the enrichment. **D)** Expression of the retinoic acid receptor *Rara* across development. Related factors such as *Rxra* show a similar pattern. **E)** Chromatin activity scores from chromVAR for the genome-wide binding sites (TFBS) of RARA and RXRA. Positive/negative scores indicate higher/lower accessibility than background chromatin. **F)** Zoomed-in region of the heatmap in B, showing the expression of genes in the cholesterol biosynthesis pathway. **G)** Same as E but for FOXC1. **H-I)** MYOD1 chromatin activity at its TFBSs (H) and gene expression (I) across stages. **J-K)** *Smad5* gene expression (J) and chromatin activity at its TFBSs (K) across stages. **L)** Gene expression levels for *Mef2c* across development. **M)** Significance scores for enrichment of chromatin regions associated with genes that show skeletal abnormalities in KO mice. The significance of each term is shown separately for the sets of regions with highest activity at each vertebral fate, indicated by the colour of the circle. Circle size is proportional to the significance level.

When ordered by developmental stage, somites from different vertebral fates showed waves of temporally restricted transcriptional and chromatin remodelling (Figure 3B, S5A). We analysed these patterns of coordinated gene expression by performing enrichment analysis of Gene Ontology functional terms. Differentially expressed genes with highest expression in cervical somites (clusters 1-4) were related to epithelial cell development and response to retinoic acid signalling, including several retinoic acid receptors (Figure 3B-C). Additional genes involved in somitogenesis and embryonic patterning were prevalent in both cervical and thoracic somites (clusters 1-6, Figure 3C), and were generally expressed at the highest level in the most-recently segmented somite. This suggests somites at these early developmental stages closely resemble their PSM lineage and are only beginning to activate a somite-specific transcriptional profile.

We used our data to dissect the complex interplay between metabolite production, TF activity and chromatin dynamics involved in retinoic acid (RA) signalling, which requires precise spatiotemporal regulation for adequate differentiation of progenitor cells (Draut, Liebenstein, and Begemann 2019). RA signalling effects are mediated by the retinoic acid receptor (RAR) and retinoid X receptor (RXR) families. These ligand-dependent transcription factors can recruit either corepressors or coactivators to induce changes in chromatin condensation and regulate transcription (Draut, Liebenstein, and Begemann 2019). Expression of the enzymes involved in RA production (*Aldh1a2* and *Rdh10*) as well as of several RAR/RXR TFs peaked early in development (Figure 3B,D). However, the accessibility of loci with binding sites for RAR/RXR factors was lowest at this stage (Figure 3E), suggesting their association with corepressors to induce chromatin condensation. As development proceeded these chromatin loci progressively increased in accessibility, maintaining an open-chromatin configuration from stage 25 onwards (Figure 3E). These data indicate that the epigenetic profile of somites is reshaped across development from a repressive to a permissive state for RA signalling activity.

Thoracic somites showed strong enrichment for genes involved in the development of the skeletal system, including both the muscle and cartilage lineages (clusters 5-6; Figure 3C). We observed prominent expression of many components of the TGFbeta, BMP and Smad signalling pathways, which are fundamental in orchestrating skeletal system development (Wu, Chen, and Li 2016) (Figure 3B). We also observed coordinated expression of cholesterol biosynthesis, with maximal expression of 17 metabolically central genes at stage 25 before being downregulated at later stages (Figure 3F). Cholesterol plays important roles in the transduction of hedgehog signalling (S. Xu and Tang 2022; Stottmann et al. 2011) and is required for the correct development of muscle and bone (Campos et al. 2015; Anderson et al. 2020). Sonic hedgehog (SHH) is secreted by the notochord and controls the specification of the sclerotome during patterning of epithelial somites (Murtaugh, Chyung, and Lassar 1999). Defective cholesterol biosynthesis leads to impaired response to Shh signalling and skeletal defects (Stottmann et al. 2011; S. Xu and Tang 2022). As expected, we did not detect significant *Shh* expression in the somites. However, the tightly controlled expression of the cholesterol pathway components suggests a mechanism to control when somites are most responsive to extrinsic hedgehog signalling.

### Somite differentiation accelerates across development

Our data revealed that chromatin remodelling is concentrated in thoracic somites. In addition to activating the gene programmes controlling skeletal system development, somites at stages 18 to 25 generally showed higher levels of open chromatin, compared to other stages. We observed a sharp increase in the fraction of reads in peaks in somites from stage 18 embryos, with further increases at stages 21 and 25, before dropping in the stage 27 somites (Figure 3G). Consistently, over half of all differentially accessible chromatin loci showed highest accessibility in thoracic somites (Figure S5A). These chromatin loci were enriched for many transcription factor motifs, including several with well described roles in skeletal system development such as forkhead TFs, implicated in both skeletal muscle and cartilage development (Sanchez, Candau, and Bernardi 2014; J. Xu et al. 2021) (Figure 3H). We also observed increased accessibility at loci harbouring binding sites for MYOD1 (Figure 3I) and MYF5 (Figure S5C), which are essential for cell commitment to the myogenic lineage (Chal and Pourquié 2017).

Previous work (Borman and Yorde 1994; Berti et al. 2015; Gi Fay Mok, Mohammed, and Sweetman 2015; Maschner et al. 2016) have characterised the expression dynamics of sclerotome and myotome markers, including *Myod1*, at several embryonic stages. These studies showed that somites from younger embryos take longer to activate marker gene expression compared to somites from more advanced embryos. We hypothesised that the shorter times required for marker expression onset in late development stem from a change in the permissiveness of the chromatin landscape, which allows lineage-defining TFs to activate their downstream pathways sooner. Analysis of the active biological processes prevalent at later developmental stages indeed revealed a switch from cell development and morphogenesis programmes to lineage commitment and differentiation (Figure 3C). These transitions were often accompanied by shifts in the active components of canonical signalling pathways, both by altering the expression of key TFs and by changing the permissiveness of the chromatin at their effector sites throughout the genome. For example, while expression of *Smad3* and *Bmp3* was at its highest levels in thoracic somites, increasing expression of *Smad5* alongside *Bmp1, Bmp4* and *Bmp7* was observed later in development (Figure 3B). *Smad5* was expressed at all stages, albeit at lower levels early on (Figure 3J); however, its binding sites only became accessible from stage 25 onwards, when expression was highest (Figure 3K). Signalling through SMAD2/3 and SMAD1/5 have opposing effects on differentiation. For example, while BMP3-SMAD3 block osteogenesis, BMP1 and BMP7, downstream of TGFbeta, promote osteoblast production (Wu, Chen, and Li 2016). Thus, the switch in usage of the opposing arms of the SMAD-BMP or SMAD-TGFbeta signalling pathways suggests cells at later developmental stages have progressed further in their differentiation trajectory. Consistent with this, genes crucial for fate determination and commitment also increased in expression across time: *Mef2c* (Figure 3L), which is fundamental in myogenic differentiation, and *Sox9* (Figure S5D), which specifies chondrocytes, peaked in sacral somites (Green et al. 2015; Molkentin et al. 1995).

Additionally, we observed significant upregulation of *Bmp4* specifically in lumbar somites (cluster7, Figure 3B). Besides regulating the development of the skeletal system, BMP4 has been shown to induce the expression of *Flk1* (*Kdr*) in epithelial somites (Nimmagadda et al. 2005), a factor essential for vasculogenesis and angiogenesis. The dermomyotome derivative lineages include vascular endothelial cells. Concomitant with *Bmp4* upregulation, we also observed increased expression of many other genes involved in angiogenesis, such as *Gata2, Tal1, Ets1* and *Tbx20* (cluster7, Figure 3B). Later in development, sacral somites continued to express high levels of angiogenic factors and further activated more mature programmes involved in blood vessel remodelling (Figure 3C).

Finally, to assess whether our molecular atlas captures regulators important in determining vertebral fate identity, we tested for enrichment of genes with specific mouse knock-out phenotypes. Genomic loci with highest activity in cervical or thoracic somites were strongly enriched for phenotypes affecting cervical vertebrae and thoracic and rib morphology. In contrast, regions active in sacral somites were associated with abnormal lumbar and sacral vertebrae (Figure 3M). Thus, our catalogue of differential activity along the axial skeleton can be utilised to identify genes and regulatory elements important for the specification of the different vertebral structures.

### Skeletogenesis is shaped by opposing regulatory modules

Next, we leveraged the paired design of our dataset to map chromatin-transcription regulatory interactions driving somite maturation and differentiation, by applying the Functional Inference of Gene Regulation (FigR) method (Kartha et al. 2022) (with minor modifications, see Methods for details). First, we computed the correlation between the activity levels of all differentially expressed genes and the open chromatin peaks within 100 kb to identify regulatory elements likely to direct nearby gene expression changes. Peak-gene pairs were considered significantly associated if they had stronger correlation values compared to randomised interactions. After restricting results to pairs with moderate to high correlation scores (>0.3), we identified 12,803 putative regulatory interactions, involving 47% of all differentially expressed genes. Although a small fraction of the interactions included promoter-like peaks in the immediate vicinity of the genes, most resembled enhancers and were dozens of kilobases away, with a median distance of 38 kb (interquartile range: 13.1-67.5 kb). Linked peaks overlapped more often with FANTOM5 and ENCODE enhancers (H3K27ac-high H3K4me3-low signature) compared to all peaks, lending support for their regulatory activity (Figure 4A). This proportion sharply increased when links were restricted to those with the strongest correlations (Figure 4A). Thus, this strategy serves to enrich the set of chromatin loci for enhancer elements, and to associate their activity to dynamically regulated genes.

**Figure 4.**
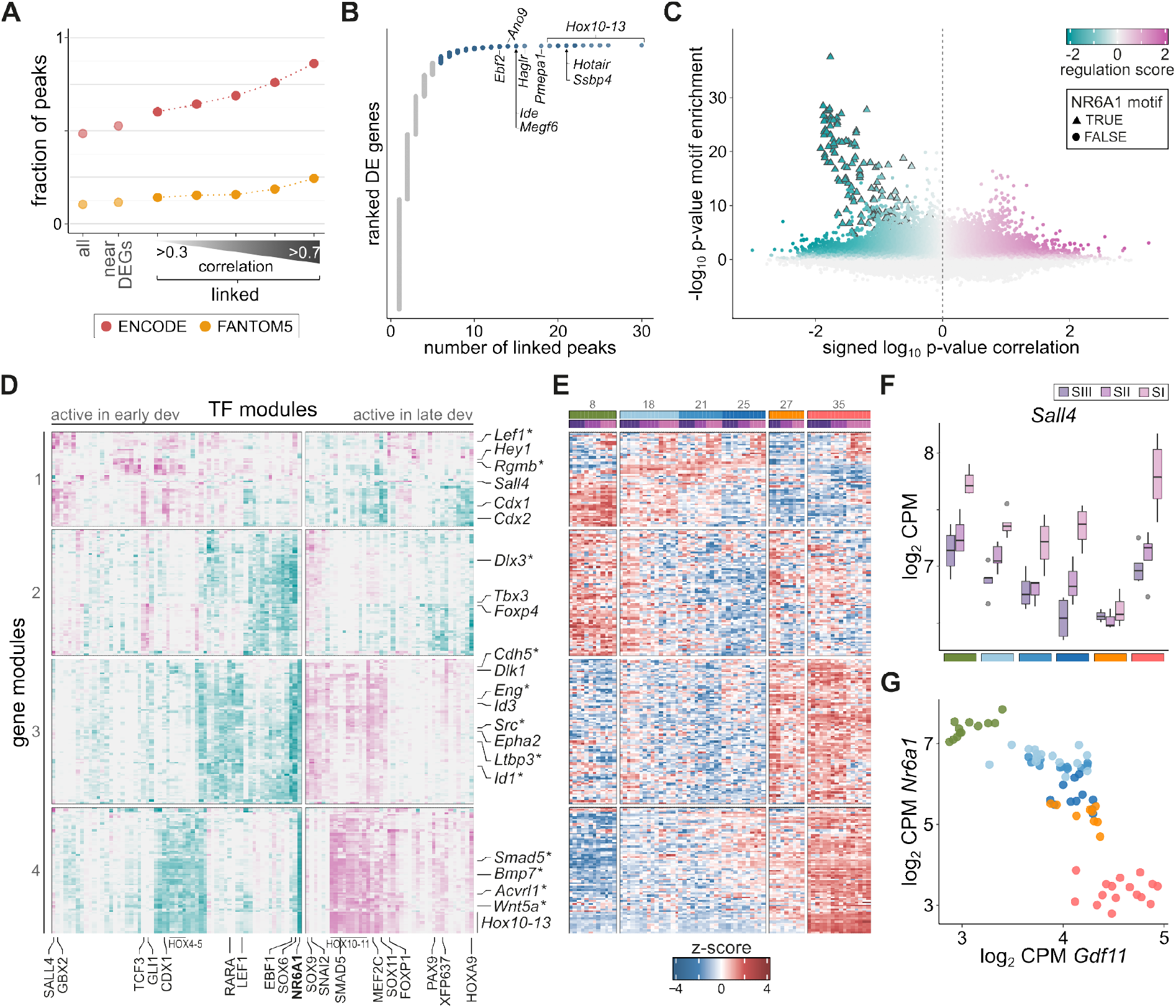
Regulatory modules with opposing activity along embryonic development ensure timely activation of skeletogenesis pathways. **A)** Fraction of peaks that overlap enhancer elements from the ENCODE and FANTOM5 catalogues. Peaks identified as putative regulators of differentially expressed genes (linked peaks) are more likely to be annotated enhancers. This fraction increases as the set of peaks is restricted to stronger interactions, as shown by limiting to linked peaks with correlation scores higher than 0.3-0.7. **B)** For each differentially expressed (DE) gene, the number of significantly associated peaks within 100kb. Several hundred genes are linked to a large number of peaks and these include many late Hox genes. **C)** Regulation scores predicted by FigR between transcription factors (TFs) and genes with many linked peaks (blue set from B). The x-axis indicates the strength of the correlation between TF expression and peak accessibility; the y-axis corresponds to the significance of the enrichment of the TF binding sites in the linked peaks. Interactions involving NR6A1 are highlighted with triangles. **D)** Heatmap depicting patterns of regulatory activity between TFs (columns) and genes (rows). Genes are split into four modules by hierarchical clustering. Genes from the TGFbeta and BMP signalling pathways are highlighted with asterisks. Colour scale is the same as in C. **E)** Heatmap showing the expression levels of the same genes as in D across all somites profiled in this study. Samples (columns) are ordered based on their observed somite stage (indicated at the top). **F)** Expression levels of *Sall4*, one of the TFs with large regulation scores on module 1 genes. **G)** Expression levels of *Nr6a1* and *Gdf11* in all somites show a strong antagonistic relationship.

Most (93.7%) differentially expressed genes significantly linked with chromatin changes were associated with one to five putative enhancer regions but a few hundred genes were linked to many more enhancers (Figure 4B). The set of 349 strongly-connected genes contained factors key in controlling somite development and differentiation, suggesting that these processes are under complex regulatory control. Some of the most highly connected genes were Hox factors from late paralogous groups (Figure 4B), consistent with chromatin remodelling playing a crucial role in controlling their timely expression (Soshnikova and Duboule 2009). Next, we scanned the peaks associated with these highly-regulated genes and identified enriched TF motifs. TF-gene pairs were assigned a *regulation score* that favours TFs showing correlated expression to the accessibility dynamics of linked peaks (Figure 4C).

We identified opposing transcriptional programmes active in early and late development by clustering transcription factors with large regulation scores (Figure 4D-E). Four different modules of regulatory activity were evident. Module 1 acted on genes that show differences in expression between the somite trios, including genes involved in the establishment of anterior-posterior patterning and somitogenesis, such as *Cdx1/2, Gbx2, Lef1*, and Hox genes from early paralogous groups (Figure 4D,E). CDX1 and GBX2 themselves, together with SALL4, showed some of the strongest regulation scores on these genes. Consistent with their role in the specification and patterning of somitic mesoderm, their expression was highest in the most immature somite I (Figure 4F).

The other three modules were instead related to genes that are downregulated (module 2) or upregulated (modules 3 and 4) with developmental progression (Figure 4E). Module 2 activity was influenced by Shh signalling (GLI1; Figure 4D), while genes expressed late in development were under the control of several transcription factors. Among these, NR6A1 showed a prominent role, particularly in module 4, with its binding sites highly enriched in the peaks linked to these genes (Figure 4C). Expression of *Nr6a1* was negatively correlated with peak accessibility, indicating a repressive regulatory effect. A number of other transcription factors, including late Hox TFs, showed large positive regulation scores on the same genes (Figure 4D), suggesting antagonistic regulatory activities to NR6A1. Among this set of activating TFs were factors key in the specification of the muscle and cartilage lineages (MEF2C and SOX9), as well as proteins involved in balancing proliferation and differentiation of the progenitor cells, with many required to avoid premature differentiation (SNAI2, ZFP637, HOXA9 and FOXP1; Figure 4D).

Recently, NR6A1 was shown to be a key regulator of the trunk-to-tail transition in the tailbud, where its expression early in development prevents premature activation of late-expressing genes, including late Hox genes. *Nr6a1* expression is then terminated by *Gdf11* to allow the trunk-to-tail transition (Chang et al. 2022). Although these regulatory interactions were dissected in undifferentiated tailbud mesoderm cells, we observed the antagonistic expression between *Nr6a1* and *Gdf11* is retained in segmented somites (Figure 4G), suggesting this regulatory programme remains at play as cells commit to the somitic lineage. Consistently, FigR predicted the strongest effects exerted by NR6A1 to affect all Hox genes in paralogous groups 10 to 13 (Figure 4D). Additional predicted regulatory interactions included several components of the TGFbeta and BMP signalling pathways from modules 3 and 4 (highlighted with asterisks in Figure 4D), with the peaks associated with these genes also showing significant enrichment for NR6A1 binding motifs. Among these genes was *Smad5*, with a regulation score only slightly lower than those observed for late Hox genes. Further, SMAD5 itself was identified as a positive regulator of module 4 genes (Figure 4D). Thus, these data suggest that the regulatory network controlled by NR6A1 not only regulates the trunk-to-tail transition in paraxial mesoderm, but it may participate in the timely activation of differentiation pathways required for skeletogenesis. As *Nr6a1* expression diminishes in later development, so does its repression of TGFbeta and BMP signalling. In turn, other transcription factors increase in activity to enhance these pathways and drive the commitment and differentiation of cells down the various somitic derivative lineages.

## DISCUSSION

The establishment of the vertebrate body plan through somitogenesis is deeply conserved. Although the molecular mechanisms driving the segmentation process are shared, alterations to the number and class of segments between different species allows facile generation of vastly different body structures among vertebrates (Gomez et al. 2008). Previous studies have characterised the transcriptional changes accompanying the transition from unsegmented mesoderm to nascent and differentiated somites in the chick (G. F. Mok et al. 2021), mouse (Chal et al. 2015) and human embryos (Xi et al. 2017), at a single developmental stage. These studies have provided insights into the molecular pathways controlling segmentation and the subsequent differentiation of somitic mesoderm derivatives.

Here, we have experimentally analysed how the three most recently segmented somites of mouse embryos remodel their transcriptional and chromatin landscapes at six different developmental stages, capturing the earliest molecular mechanisms that give rise to cervical, thoracic, lumbar and sacral structures. We identified three thousand genes transcriptionally remodelled during somite maturation. The expression of these genes after segmentation follows the same pattern at independent stages, indicating that somites from different axial levels adhere to a conserved differentiation trajectory. However, we also identified genes expressed in specific developmental stages of the embryo, reflecting changes in the microenvironment in which somites develop.

Although the pace with which new somites are formed is roughly constant across development (Palmeirim et al. 1997), our data reveals that somite differentiation accelerates as embryos grow. We observed that somites from stage 8 embryos maintain a naive transcriptional profile for the entirety of the 6 hours captured in our data, but progression along development results in a shortening of the time spent in such an undifferentiated state. At later stages, somitic gene regulation is dominated by transcription factors controlling cell fate commitment. By the time embryos have formed 35 pairs of somites, differentiation programmes of derivative lineages are upregulated within a few hours post-segmentation and can already be observed in somite III. This is consistent with previous work in chick embryos (Borman and Yorde 1994; Berti et al. 2015; Gi Fay Mok, Mohammed, and Sweetman 2015; Maschner et al. 2016) that showed pronounced differences in the onset of expression of key factors for the commitment of cells to the myogenic lineage, depending on embryonic age. Our data characterises the regulatory mechanisms controlling cell fate specification and commitment well before the onset of definitive lineage markers, and revealed that the acceleration of somite differentiation in later development initiates soon after somite segmentation. This phenomenon is not restricted to the myogenic lineage, but extends to other cell type populations, including chondrogenic and endothelial cells. In sum, our high-resolution view of the molecular mechanisms underlying the specification and development of somitic lineages has revealed novel features of somitogenesis and is a powerful resource for the developmental biology community to study its progression in mammals.

## METHODS

### Mouse embryo collection and dissection

All experiments followed the Animals (Scientific Procedures) Act 1986 (United Kingdom) and with the approval of the Cancer Research UK Cambridge Institute Animal Welfare and Ethical Review Body (form number: NRWF-DO-01-v3). Animal experiments conformed to the Animal Research: Reporting of *In Vivo* Experiments (ARRIVE) guidelines developed by the National Centre for the Replacement, Refinement and Reduction of Animals in Research (NC3Rs). C57BL/6J strain mice were obtained from Charles River Laboratories and maintained under standard husbandry practices.

Mouse embryos from the appropriate somite stages (8 to 35 somites) were dissected in RNAse-free conditions in cold PBS, on silicone plates. Utmost care was taken to accurately count the number of somite pairs of each embryo; however, for the 35-somite stage embryos this task becomes very difficult and it is possible that there is a one or two somite error range in the number estimated. Photos of all the embryos profiled are provided in Supplementary File 1. To dissect out the somites, embryos were treated with dispase II (1mg/mL in DMEM) for 30-45 seconds at 37°C. The three most posterior pairs of somites were then collected using tungsten needles, dissecting out every somite separately. We labelled each somite pair as somite I, II or III from the most posterior to the most anterior, respectively. Thus, somite I corresponds to the most recently segmented somite, while somites II and III were segmented ~2 and ~4 hours before (Dequéant and Pourquié 2008); we refer to this as the somite’s age. Each individual somite was placed in 10μL of lysis buffer (Takara) containing RNase inhibitor. One somite from each pair was flash frozen in liquid nitrogen and stored at −80°C for later processing for RNA-seq. The matching somites were directly processed to generate ATAC-seq libraries.

### Experimental design

We collected at least four different embryos from each developmental stage (Table S1). Dissections were performed on ten different days with every stage represented on at least two separate collection dates. However, samples from three different stages were collected on five days, without overlap with the samples from the remaining three stages, resulting in a partially confounded design.

### RNA-seq experiments and library preparation

Reverse transcription was performed directly on frozen lysed somites and cDNA was amplified with 8 cycles of PCR, using the SMART-seq v4 Ultra Low Input RNA Kit for Sequencing (TaKaRa, 634891). RNA-seq libraries were generated from 100pg of amplified cDNA using the NEXTERA XT DNA Library Preparation kit (Illumina, FC-131-1096), according to the manufacturer’s instructions, except only a quarter of the recommended reagents’ amount was used. The resulting libraries were quantified using a Qubit instrument and their size distributions were assessed with a TapeStation machine. Pooled libraries were sequenced on an Illumina HiSeq 4000 according to manufacturer’s instructions to produce paired-end 150bp reads.

### ATAC-seq experiments and library preparation

ATAC-seq experiments were performed following the protocol from Corces and colleagues (Corces et al. 2017). Briefly, individual somites were lysed and transposed with 1μL of transposome (Nextera DNA Sample Preparation kit FC-121-1030) at 37°C for 30 minutes. Samples were then purified with the Zymo Clean & Concentrator kit and eluted in 21μL of elution buffer. Transposed DNA was quantified by qPCR using 5 μl of PCR products. The number of additional cycles was determined by plotting linear Rn versus cycle and corresponded to one third of the maximum fluorescence intensity. Transposed DNA was then amplified with 13 cycles of PCR. The final products were double size-selected with AMPure beads (0.55X - 1.5X) to obtain fragments between 100bp and 700bp. Libraries were quantified and the sizes were assessed with a TapeStation machine. Pooled libraries were sequenced on an Illumina HiSeq 4000 according to manufacturer’s instructions to produce paired-end 150bp reads. Samples were sequenced to a median depth of 70.5 million fragments.

### RNA-seq data processing and quality control

RNA-seq paired-end fragments were aligned to the mouse reference genome (GRCm38) with STAR 2.6.0c (Dobin et al. 2013) with options --outFilterMismatchNmax 6 -- outFilterMatchNminOverLread 0.5 --outFilterScoreMinOverLread 0.5 --outSAMtype BAM SortedByCoordinate --outFilterType BySJout --outFilterMultimapNmax 20 --alignSJoverhangMin 8 --alignSJDBoverhangMin 1 --alignIntronMin 20 --alignIntronMax 1000000 --alignMatesGapMax 1000000 --outSAMstrandField intronMotif. On average, 84% of the sequencing fragments mapped uniquely. We also set the option --quantMode GeneCounts to quantify the number of fragments overlapping annotated transcripts, using Ensembl’s genome annotation version 96 (http://apr2019.archive.ensembl.org/index.html).

Samples were sequenced to a median depth of 17.4 million paired-end fragments. One sample had a library size of only 88 thousand fragments and was discarded. All other samples showed a uniform number of fragments mapped uniquely (median 85.6%, standard deviation (SD) 5.6%) and most of these were within annotated exons (median 84.9%, SD 2.8%). On average, we detected around 22 thousand expressed genes per sample (Table S2).

To validate the staging of samples we exploited the Hox code that serves as a molecular indicative of developmental stage. As shown in Figure 1C-D, our embryo stages defined by the observed number of somites agreed with the expected expression levels of Hox genes. However, samples from one stage 27 embryo were more similar to the stage 35 somites, showing expression of several late Hox genes from paralogous groups 12 and 13 that are only observed in the stage 35 samples. These data suggest this particular embryo was likely of a more advanced stage than 27 somites and was removed from downstream analyses (Table S1).

### RNA-seq data normalisation

Downstream analyses were restricted to genes with at least 10 counts in three or more samples, as implemented in the filterByExpr function from the edgeR package (Mark D. Robinson, McCarthy, and Smyth 2010; McCarthy, Chen, and Smyth 2012); this represents 20,062 genes. To normalise for differences in sequencing depth we used the calcNormFactors function that implements the weighted trimmed mean of M-values method (M. D. Robinson and Oshlack 2010), and generated counts-per-million normalised expression estimates.

A PCA of the thousand most variable genes (determined from variance-stabilised data, computed with the vst function form the DESeq2 package) revealed good separation of samples from different developmental stages (Figure S2A). However, we also observed subgrouping by the date of collection, indicating substantial batch effects (Figure S2A). Since the experimental design is partially confounded with the date of collection of the samples, we were unable to include this as a covariate in downstream analyses. Instead, to control for technical variation unrelated to the biological variables of interest, we used the function lmFit from the limma package (Ritchie et al. 2015) to fit a linear model of the combination of developmental stage and somite age for each sample. We then performed PCA on the residuals from the fit (function residuals) to capture systematic variation unrelated to the biological design of interest. To determine how many principal components (PCs) captured significant variation we used the parallelPCA function from the scran package (Lun, McCarthy, and Marioni 2016) on the normalised counts; this function estimates, via permutation analysis, the number of PCs that explain more variation than expected by chance, which in our case was 14. Thus, the 14 first PCs were used as covariates in downstream analyses to control for unwanted variation (Figure S2B). We note that this procedure captures both technical and biological variation not modelled in our design of interest (i.e. the sex of the embryos).

### RNA-seq differential expression analysis

To identify genes significantly differentially expressed across conditions we used edgeR (Mark D. Robinson, McCarthy, and Smyth 2010; McCarthy, Chen, and Smyth 2012), with a design matrix of the interaction of each sample’s age and developmental stage, plus the 14 PCs representing technical variation as covariates. Dispersion was estimated with the estimateDisp function (setting robust = TRUE) and fitting the model with glmQLFit. Specific contrasts were then tested with the glmQLFTest function.

To identify the regions that change as somites differentiate we compared samples from somites I, II and III. To identify conserved differences across development we defined contrasts for all three pairwise comparisons, using the average of same-age samples from all six stages:

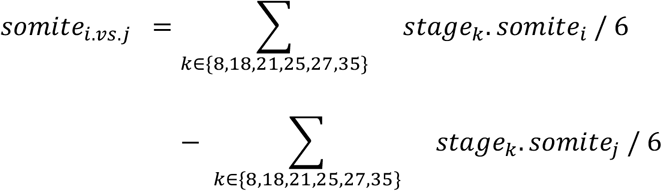

where *i.vs.j* corresponds to *I.vs.II, I.vs.III* and *II.vs.III*. To recover possible stage-specific changes, we also defined contrasts on a per-stage basis:

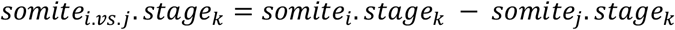

where *k* is one of the six stages and *i.vs.j* the same as above. All three pairwise comparisons from each stage were tested at once. Thus, in these cases the p-value indicates whether the gene is differentially expressed between at least a pair of somite ages.

We used a similar approach to test for differences in expression across development. Conserved differences between all somites irrespective of their maturity level were assessed by averaging somites I, II and III and testing each pairwise comparison between the six stages:

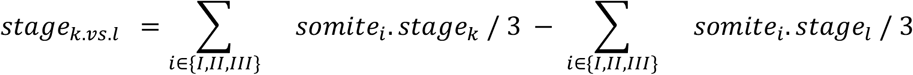

where *k.vs.l* corresponds to all pairwise comparisons between the six stages. All contrasts were tested at once to avoid performing too many tests and, again, p-values indicate whether the gene is significantly different between at least a pair of stages. To check for any changes specific to a given somite age we repeated the analysis separately for somites I, II and III:

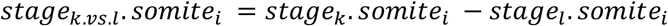

where *i* is any of the three somite ages and *k.vs.l* the same as above. Genes were considered significantly differentially expressed if their adjusted p-value was lower than 0.05 (FDR < 5%) and their absolute fold-change was greater than 1.5. Results from all differential expression analyses are available at https://github.com/xibarrasoria/somitogenesis2022.

### ATAC-seq data alignment

Raw sequencing reads were aligned to the mouse reference genome (GRCm38) using bwa mem 0.7.12-r1039 (Li and Durbin 2009) with default parameters. On average, 93% of the total fragments were successfully aligned. The resulting SAM files were processed with samtools 1.5 (Li et al. 2009). One sample was sequenced to a disproportionately high depth compared to the rest (1.5 billion fragments compared to a median of 70.5 million). The mapped data for this sample was downsampled to 15% of the total reads (samtools -s 0.15), and the resulting BAM file was used in the downstream processing steps.

Duplicated fragments were marked and removed using MarkDuplicates 1.103 from Picard tools (http://broadinstitute.github.io/picard) with option REMOVE_DUPLICATES=TRUE. We further used samtools to remove any pairs that were not properly aligned (-f 0×02); supplementary alignments (-F 0×800); alignments with mapping quality lower than 30 (-q 30); and alignments outside the autosomes or chromosome X. The resulting BAM files represent the clean, good quality alignments used in all downstream analyses.

### ATAC-seq quality control

To assess the quality of the libraries we used three different criteria: 1) the insert size distribution of the sequenced fragments; 2) the level of signal enrichment at the transcription start site (TSS) of expressed genes; and 3) the signal-to-noise ratio, assessed by the ability to call peaks (Figure S3 and Table S3).

To compute the insert size distribution of each library we used the getPESizes function from the csaw package (Lun and Smyth 2016). Each library’s distribution was visually inspected and scored based on the number of nucleosomal peaks. Thus, a score of 0 implies that only short fragments were recovered, a score of 1 indicates presence of mononucleosomes, 2 corresponds to samples with both monomers and dimers, and so on. The maximum score assigned was 4, including samples with fragment sizes corresponding to nucleosome tetramers or larger (Figure S3A).

To estimate the enrichment of fragments at transcription start sites we applied the method recommended by the ENCODE standards for ATAC-seq data (https://www.encodeproject.org/data-standards/terms/#enrichment). Specifically, we restricted the analysis to genes with moderate to high expression as assessed from the RNA-seq data (mean normalised counts per million greater than 10, corresponding to 8,025 genes). We then used the biomaRt package (Durinck et al. 2005) to extract the most 5’ TSS for each gene and created a BED file of 2kb intervals centred at each TSS. We computed the coverage of such intervals using bedtools coverage 2.26.0 (Quinlan and Hall 2010) a BEDPE file containing the Tn5 insertion sites inferred from the aligned fragments (by shifting the start/end coordinates by +5/-4 bp with an ad hoc perl script). To calculate the enrichment at the TSS we first computed the mean insertion counts at each base pair from all genes. We then used the mean of the first and last 100bp as an estimate of the background insertion rate. For each base pair, we computed the enrichment score as the fold-change against the background rate; this results in an enrichment score of ~1 at the flanks of the 2kb interval which increases as it approaches the TSS (Figure S3B).

Finally, to assess the signal-to-noise ratio of each sample we used MACS2 2.1 (Zhang et al. 2008) to call peaks, with options callpeak -f BAMPE -g mm --keep-dup all --broad. Peaks overlapping blacklisted regions (Amemiya, Kundaje, and Boyle 2019) (obtained from https://github.com/Boyle-Lab/Blacklist/blob/master/lists/mm10-blacklist.v2.bed.gz) were discarded. We calculated the fraction of reads in peaks (FRiP) as the total fragments overlapping called peaks over the total library size and used this, along with the total number of peaks, as proxies for the signal-to-noise ratio (Figure S3C).

Libraries with an insert size distribution showing a good nucleosomal pattern generally had good TSS enrichment scores and signal-to-noise ratios. For each sample we defined a quality control pass if they had an insert size distribution score of 2 or higher; a fraction of reads in peaks of 3% or larger; at least 15,000 peaks; and a TSS enrichment score of 5 or higher (Figure S3D). Samples satisfying at least three of these criteria were annotated as good quality and used in downstream analyses (50 of the 75 libraries). Importantly, insert size distribution scores were positively correlated with the experimentally measured DNA fragment sizes but showed no relation to sequencing depth, indicating that samples with poor quality control characteristics are not due to insufficient sequencing (Figure S3E).

### ATAC-seq peak calling

To define a unified set of peaks for the whole dataset we combined the clean BAM files from the 50 samples that passed quality control and used them as input for MACS2 (same parameters as stated above). By calling peaks on the combined data from all samples, the peak calling process becomes agnostic to the different conditions in our experimental design, which is important for downstream differential accessibility analyses. After removing peaks overlapping blacklisted regions, a total of 131,743 peaks were called, with a median width of 777 bp (interquartile range 418-1394 bp). We re-computed the FRiP for each sample using this common peak set.

It is possible that by merging all samples together, some low-enrichment stage-specific peaks are lost. Thus, we repeated the peak calling procedure but on a stage-specific basis. When comparing the per-stage peak calls to the set obtained by using all 50 samples, around 96.6% of the peaks called in each stage were also called in the all-sample set (range 93.43-98.09%). The small proportion of peaks missed generally had small fold-changes and high q-values, and thus correspond to low significance calls. This provides confidence that we have not missed stage-specific peaks by merging data from all samples.

### ATAC-seq data normalisation

To normalise the ATAC-seq data we used the methods implemented in the csaw package (Lun and Smyth 2016). We generated MA plots comparing all pairs of samples by counting the number of fragments in 10 kb windows tiling the genome. For high-abundance windows, which correspond to open chromatin, we observed a deviation of the log_2_ fold-change from the expected value of 0 that was correlated with the abundance level of the genomic region (Figure S4A). Conventional normalisation techniques used in the majority of ATAC-seq analyses compute a single size factor that captures systematic differences between samples; these approaches fail to account for the trend observed in our data (Figure S4B). Thus, we instead used a loess-based approach to compute size factors specific to each abundance level. For this, we counted the number of fragments mapped to 150 bp windows, sliding along the genome by 50 bp, using the function windowCounts (with filter set to 75 and excluding any reads overlapping blacklisted regions). We then filtered out any windows that did not overlap the common peak set or that had less than an average count of 4 fragments across samples. We finally used this set of windows to compute the size factors with the normOffsets function (with type=loess). This approach successfully removed the observed trend (Figure S4C). However, the first principal component estimated from the normalised counts of the 5000 most variable windows was strongly correlated with samples’ FRiP (Pearson’s r = −0.59), suggesting other technical effects were still dominant in the data (Figure S4D).

To remove unwanted variation from the dataset we used the same strategy that we applied to the RNA-seq samples. That is, we obtained the residuals from a linear model fit of the interaction of the stage and somite age of each sample and applied PCA to capture the major sources of variation. We retained the first 18 PCs, since these were deemed to explain significantly more variation than chance (as determined by the parallelPCA function), which significantly removed efficiency and batch effects (Figure S4E).

### ATAC-seq differential accessibility analysis

To test for differences in accessibility across conditions we used the approach implemented in csaw (Lun and Smyth 2016). We based the analysis on the window counts described above. Window counts along with the corresponding size factors were converted into a DGEList object compatible with edgeR (Mark D. Robinson, McCarthy, and Smyth 2010) to perform differential analysis. The same approach as described for the RNA-seq data was used.

After each window was tested, we used the mergeWindows function to merge windows that were no more than 150 bp apart, restricting the maximum width to 1.5 kb. Regions larger than 1.5 kb were broken into smaller overlapping regions of roughly equal size (+/- 100bp). We then computed a combined p-value for each of these regions with the combineTests function, using Simes’ method. Correction for multiple testing was performed at the region level and regions were considered significantly different if their adjusted p-value was lower than 0.05 and their absolute fold-change was greater than 1.5. Results from all differential expression analyses are available at https://github.com/xibarrasoria/somitogenesis2022.

### Functional terms enrichment analysis

Gene ontology enrichment analysis was performed using the elim method from the TopGo package (Alexa and Rahnenfuhrer 2022), as implemented in the topGOtable function from the PCAexplorer 2.18.0 package (Marini and Binder 2019). Enrichment of GO terms among differentially expressed genes was computed, using all genes expressed in somites as the background.

Enrichment analysis of GO terms and mouse KO phenotypes in differentially accessible chromatin regions was computed with GREAT (McLean et al. 2010), using the implementation from the rGREAT 1.24.0 package (Gu and Hübschmann 2022). All somite peaks were used as the background set.

### Motif enrichment analysis

To determine transcription factor (TF) motifs enriched in the regions of open chromatin, we used Analysis of Motif Enrichment (McLeay and Bailey 2010) from the MEME suite (Bailey et al. 2009), with the human and mouse HOCOMOCOv11_full motif databases. Enrichment in differentially accessible regions (either between somite ages, or between stages, split by the vertebral fate showing highest accessibility) was computed by comparing to a set of non-differentially accessible regions with a similar length distribution.

### chromVAR

To estimate the accessibility dynamics of sites harbouring specific TF binding sites we used chromVAR 1.12.0 (Schep et al. 2017), on the normalised and corrected window counts described above. Following the authors’ recommendations, we removed overlapping windows with the filterPeaks function, and then scanned them for matches to the motif collection provided with the package (mouse_pwms_v2). Accessibility deviation scores were then computed with the computeDeviations function. For all plots, we use the z-scores returned by this function.

### FigR

To infer peak-gene putative regulatory links we used FigR 0.1.0 (Kartha et al. 2022), restricted to the normalised and corrected counts of the 43 samples with both RNA and ATAC-seq profiles available. The function runGenePeakcorr was used to compute the correlation between each differentially expressed gene and all peaks within 100kb. This function uses chromVAR to determine a set of 100 background peaks matched for accessibility and GC content levels, to determine if the observed correlation of the gene-peak of interest is significantly higher than the correlation of the gene to these unrelated background peaks. chromVAR samples with replacement to define the background set. When there are not enough matching peaks, the number of *different* peaks in the background set can drop substantially; in extreme cases this can result in a single peak repeated 100 times. Using this distribution as a null is non-informative and thus the p-values computed for such gene-peak pairs are misleading. To avoid these cases, we modified the code to check for the number of different peaks included in the background and set the p-values to NA for any gene-peak pairs with background sets containing fewer than 50 different peaks. Downstream analyses were performed using gene-peak pairs with a p-value < 0.05 and a correlation score greater than 0.3. To infer regulatory interactions, we used the getDORCScores and runFigRGRN functions, on all genes with more than 5 linked peaks. TF-gene pairs with an absolute regulation score greater than 1.25 are considered putative interactions (Figure 4C-D).

## Data availability

The raw and processed data from this study are available in the ArrayExpress repository and can be accessed through the BioStudies database (https://www.ebi.ac.uk/biostudies/) under accession numbers E-MTAB-12511 for the RNA-seq dataset and E-MTAB-12539 for the ATAC-seq dataset. These include the raw FASTQ files as well as raw, normalised and batch-corrected count tables. All code used for data processing and analysis are available at https://github.com/xibarrasoria/somitogenesis2022.

## Acknowledgements and funding

The authors would like to thank Aaron Lun for valuable suggestions on the analytical strategy, Pascale Gilardi-Hebenstreit for insightful comments on the manuscript and CRUK Cambridge Institute Bioinformatics and Biological Resources Unit Core facilities. This work was supported by Cancer Research UK (ET, DTO: 20412, 22398), the Wellcome Trust (XIS: 108438/Z/15 to JCM; and 202878/Z/16/Z to DTO), the European Molecular Biology Laboratory (JCM), the European Research Council (DTO: 615584, 788937), and Helmholtz NCT funding (DTO: DKFZ Abteiling B270).

## Author contributions

ET, XIS, JCM, DTO designed the study; ET performed experiments; AEM, GFM provided microdissection training; XIS analysed the data; XIS, JCM, DTO wrote the manuscript; XIS, JCM, DTO oversaw the work. All authors read and approved the final manuscript.

## SUPPLEMENTAL INFORMATION

**Figure S1.**
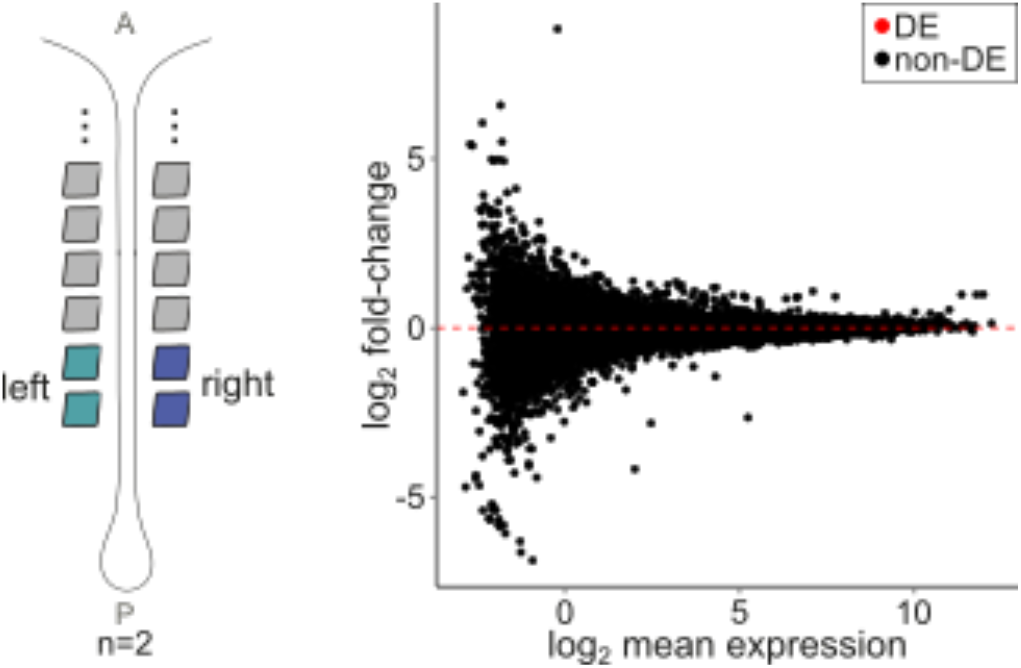
The left and right somites are transcriptionally equivalent. We compared the transcriptomes of matched left and right somites by RNA-seq using the two most posterior somite pairs, as shown in the schematic. The scatter plot shows the average gene expression level on the x-axis and the corresponding fold-change between the left and right somites on the y-axis. No significantly differentially expressed (DE) genes were identified (FDR 5%), indicating that the transcriptomes of the two somites from the same pair are equivalent. A: anterior; P: posterior.

**Figure S2.**
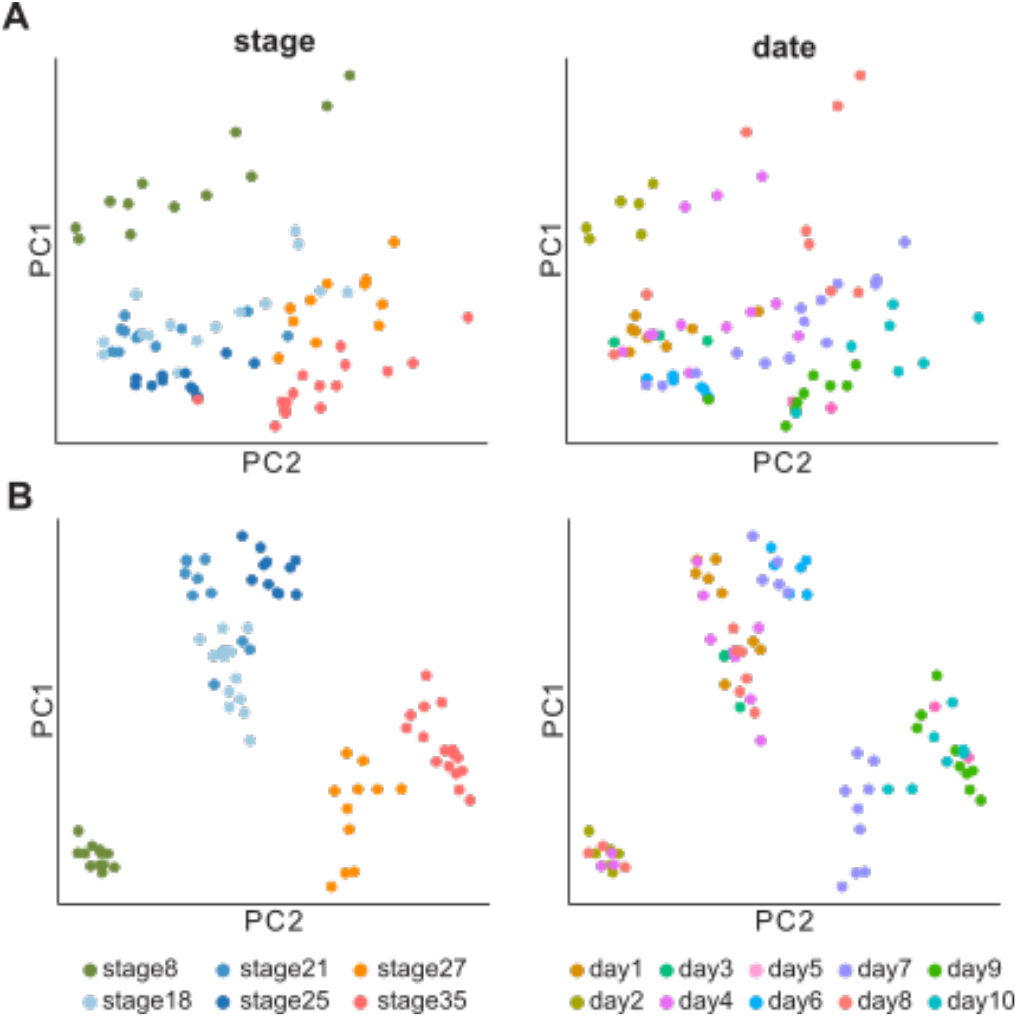
Batch correction of RNA-seq data. **A)** PCA of the normalised counts of the thousand most variable genes across samples. There is clear separation by developmental stage (left). However, PC1 also separates samples based on their collection day (right). **B)** PCA after regressing out covariates capturing variation unrelated to the experimental design. Samples separate better by their developmental stage (left) and grouping by collection day is no longer evident (right).

**Figure S3.**
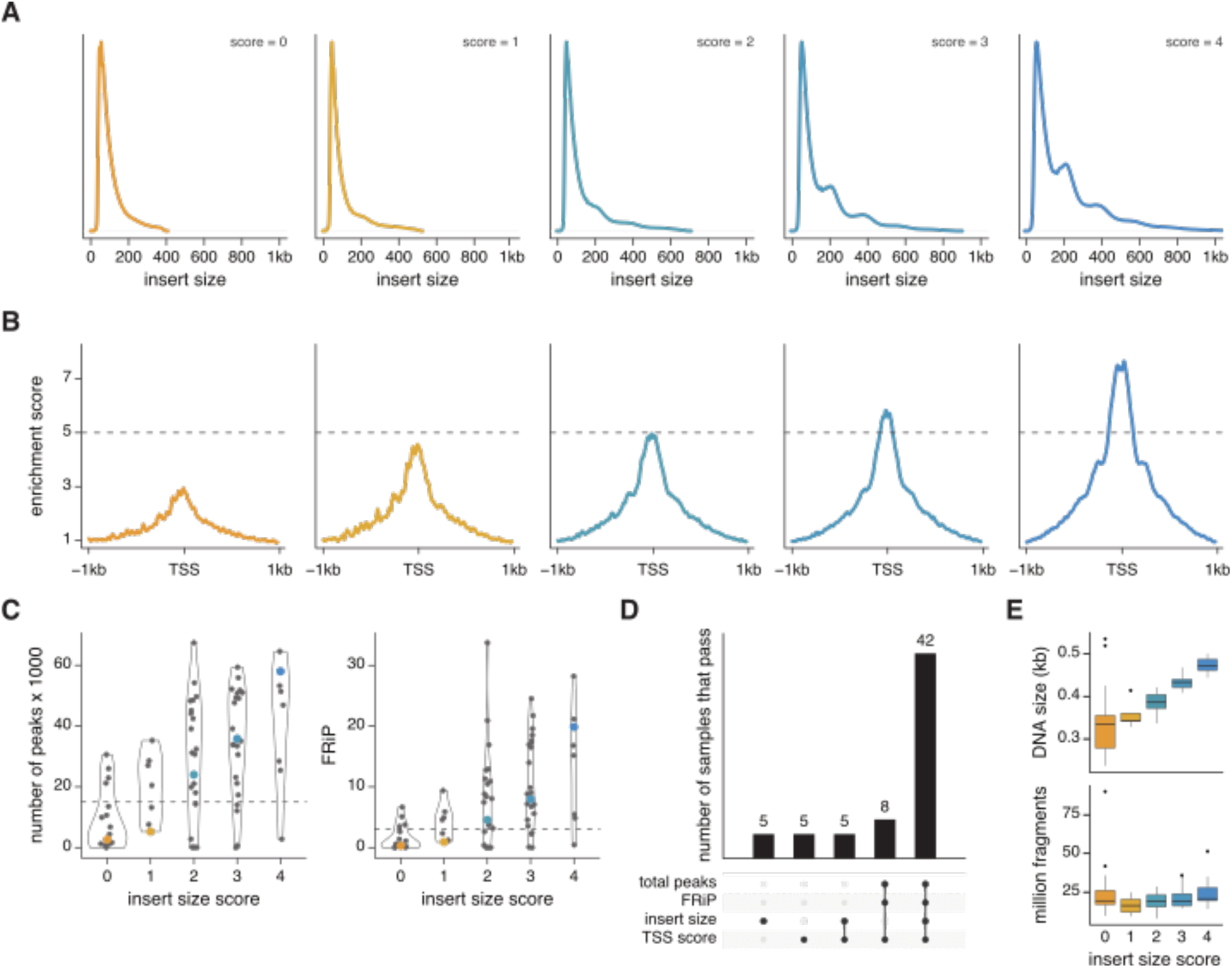
Quality control of ATAC-seq libraries. Several metrics were used to determine if the ATAC-seq libraries were of good quality. **A)** Representative density plots of the fragment sizes of the sequencing data for each score group. Scores were determined based on the number of nucleosomal peaks. **B)** Representative plots of the cumulative signal for 2 kb intervals centred at the transcription start site (TSS) of expressed genes, for the same samples shown in A. An enrichment score larger than 1 indicates an excess of insertions relative to background. Signal is smoothed by taking the rolling median of 25 bp intervals. **C)** Violin plots depicting the number of total peaks called from each library (left) and the fraction of reads in peaks (FRiP; right), stratified by the sample’s insert size distribution score. The samples depicted in A-B are highlighted by coloured points. **D)** Number of samples that pass each of the QC criteria. The 50 samples that passed three or four criteria were deemed of good enough quality for downstream analyses. Samples that failed all four criteria are not shown. **E)** Boxplots of the experimentally determined DNA fragment size (top) and the library size of the sequenced samples (bottom), stratified by the insert size distribution score. No relationship is observed between library size and size distribution score, indicating that the poor-quality samples are not due to insufficient sequencing.

**Figure S4.**
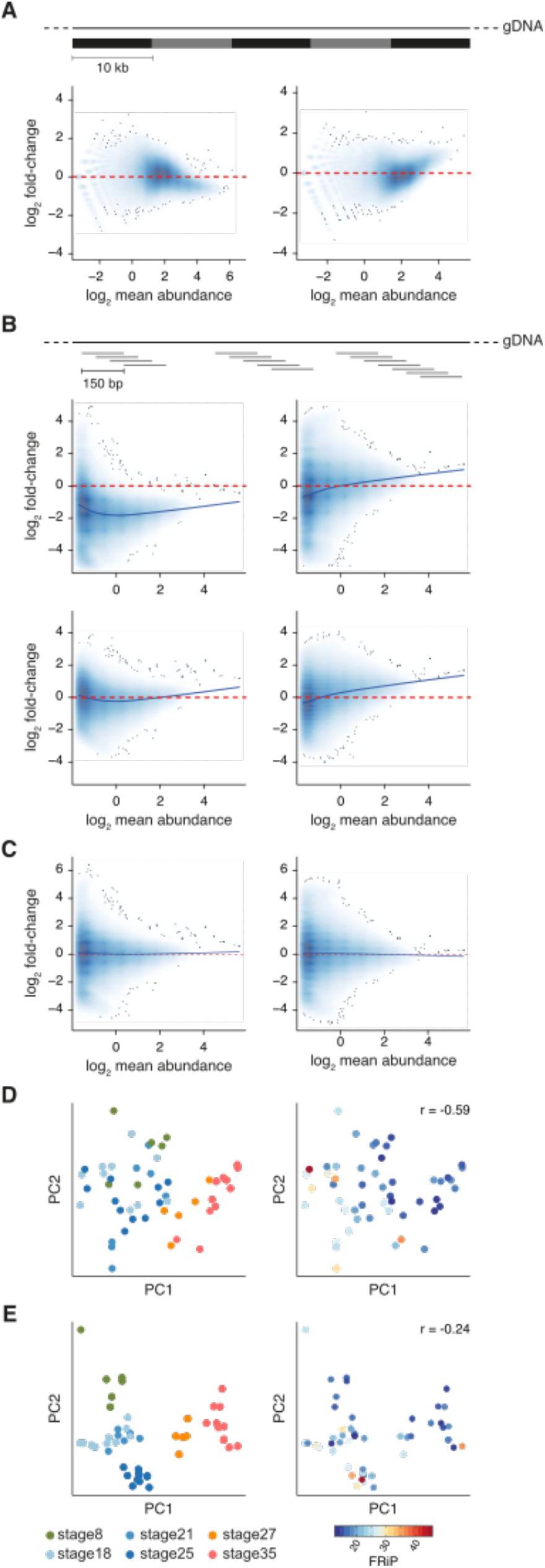
Normalisation of ATAC-seq data. **A)** MA plots of two representative samples. On the x-axis is the log_2_ average number of sequencing fragments in 10kb bins covering the mouse genome. The y-axis corresponds to the log_2_ fold-change against a reference sample (same for both panels). Comparable samples should show log fold-changes centred around 0. High abundance bins (which contain open regions) show significant deviation from 0. This deviation shows a trend dependent on mean abundance. **B)** To normalise the observed biases we focused on the regions of open chromatin. The MA plots now show on the x-axis the average counts in 150bp windows that slide 50bp, restricted to regions overlapping called peaks. At the top, the raw counts for the same samples in A. At the bottom, scaling normalisation is applied, which results in a shift towards a fold-change of 0. However, the observed trend dependent on mean abundance is still present in the data. **C)** MA plots for the same samples in B, but after applying loess-based normalisation, which computes a size factor for each abundance level. This successfully captured and removed the observed trend, with windows now centred around 0. **D)** PCA of the normalised counts of the 5000 most variable windows. On the left, samples are coloured by developmental stage while on the right they are coloured by their fraction of reads in peaks (FRiP). There is clear grouping of samples based on their FRiP. The Pearson correlation coefficient between FRiP and PC1 is noted. **E)** PCA after regressing out covariates capturing variation unrelated to the experimental design. Samples separate better by their developmental stage (left) and there is no longer a significant correlation with FRiP (right).

**Figure S5.**
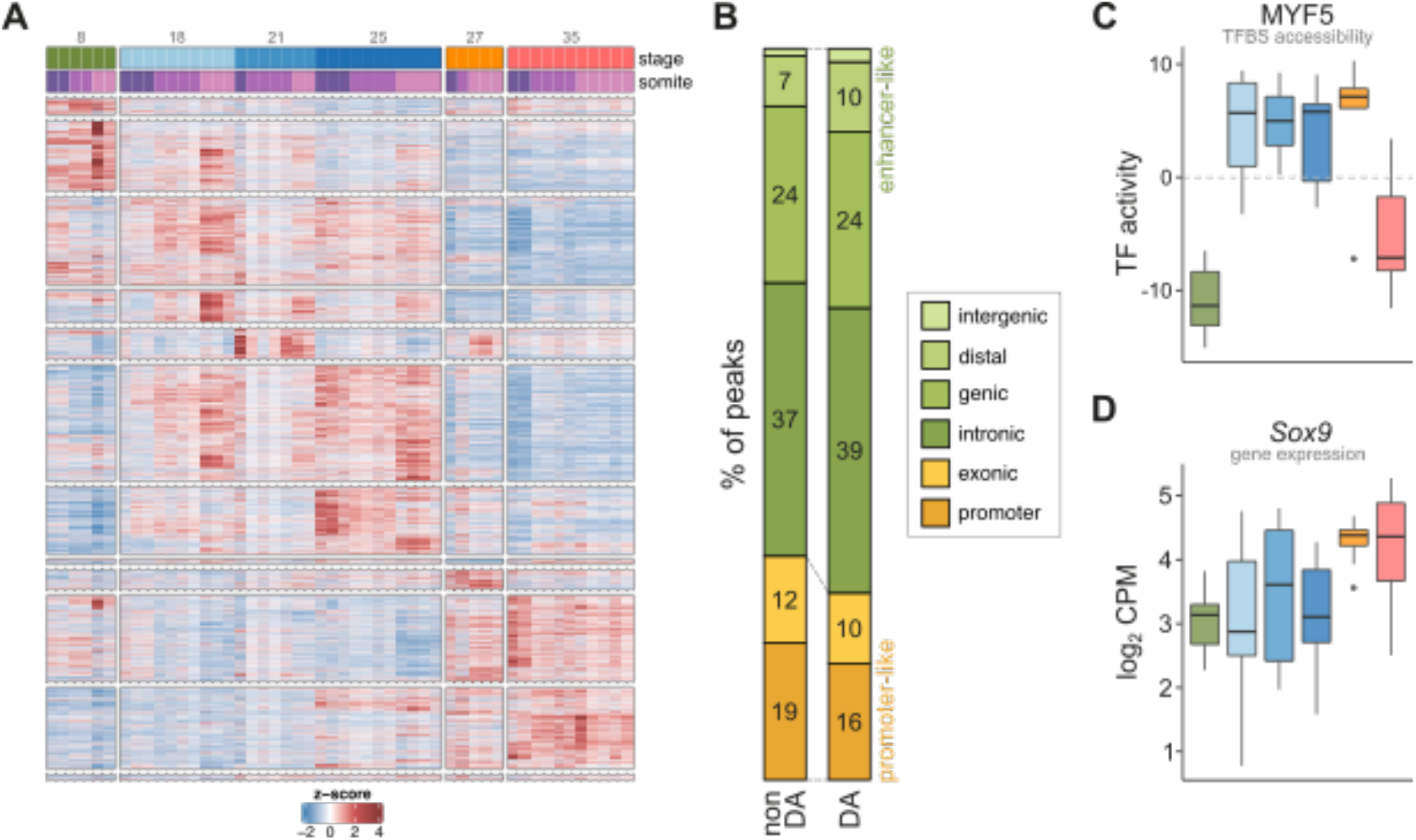
Differentially accessible chromatin loci across development. **A)** Similar to Figure 3B but showing the accessibility levels of all differential peaks across development. Samples (columns) are ordered based on their observed somite number, and their stage and somite level are indicated at the top. Peaks (rows) are split into clusters by hierarchical clustering. **B)** Same as Figure 2E but for differentially accessible regions across development. **C)** Chromatin activity scores from chromVAR for the genome-wide binding sites (TFBS) of MYF5 across development. **D)** Gene expression levels for *Sox9* across development.

**File S1 |** Photos of all the embryos profiled in this study.

**Table S1 | Metadata of the samples collected and the RNA- and ATAC-seq libraries produced.** The QC columns indicate whether the sample passed quality control (1) or not (0; highlighted in red); NA indicates that the sample did not yield a successful library for sequencing.

**Table S2 | Quality control statistics from the RNA-seq libraries.** Only one sample failed QC (in red) due to insufficient sequencing depth (totalFragments). The ‘uniqueInExons’ column indicates the number of fragments uniquely mapped to the genome that also can be assigned unambiguously to annotated exons. The ‘numberGenes’ column indicates the total number of genes with at least one count.

**Table S3 | Quality control statistics from the ATAC-seq libraries.** A third of the samples failed QC (in red). The ‘unique’ column corresponds to the number of fragments retained after removing PCR duplicates. The ‘insertSizeDist’, ‘numberPeaks’, ‘readsInPeaks’ and ‘TSSenrichment’ columns correspond to the metrics used to determine if a sample was of good quality (see Figure S3 for details).

