## Supplementary material for "A transcriptional and regulatory map of mouse somitogenesis": File_S1

### Stage: 8 somite pairs

| e5 | e6 |
| --- | --- |
| 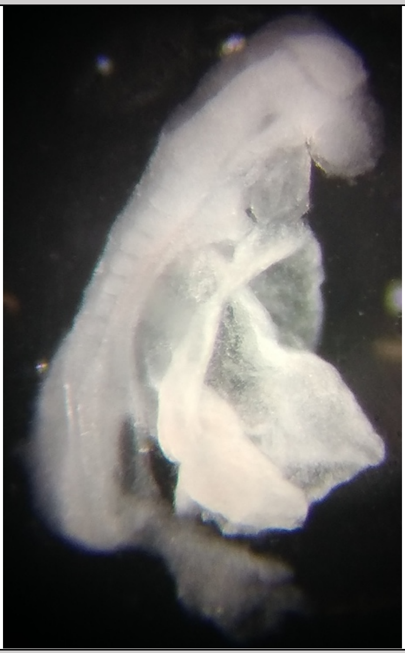 | 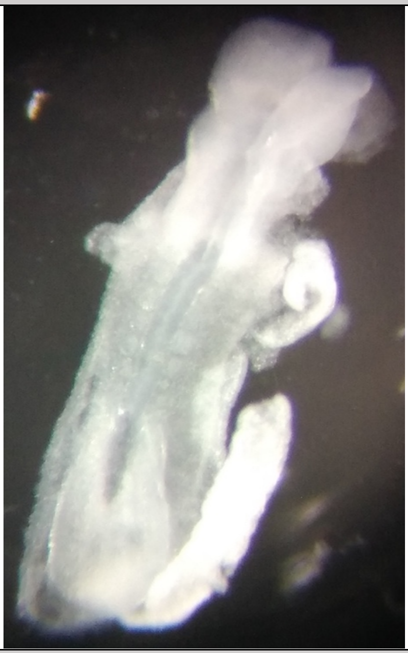 |
| **e16** | **e26** |
| 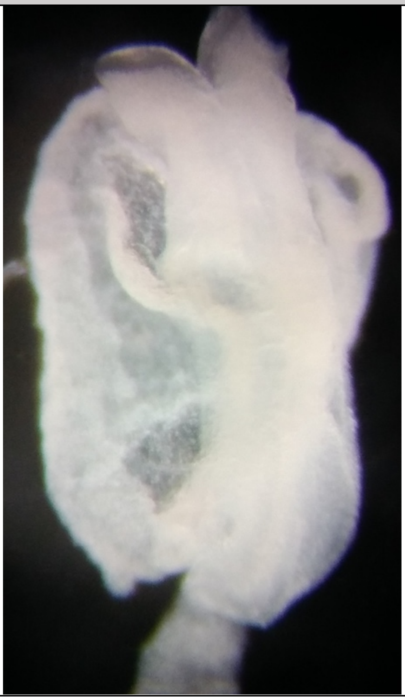 | 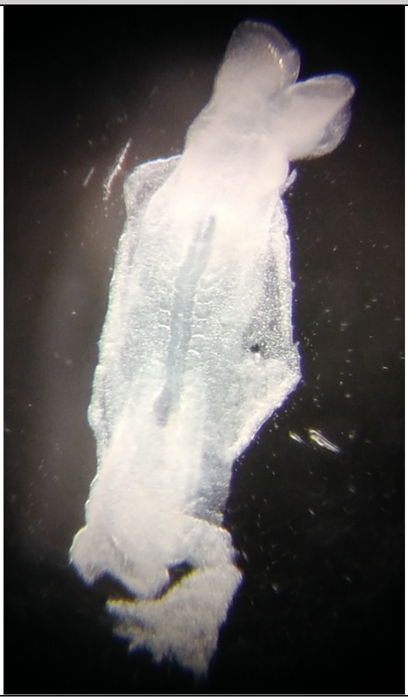 |

### Stage: 18 somite pairs

| **e9** | **e14** |
|---|---|
| 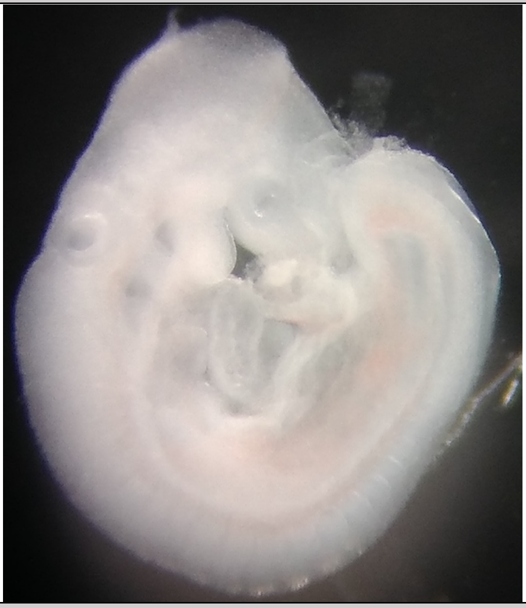 | 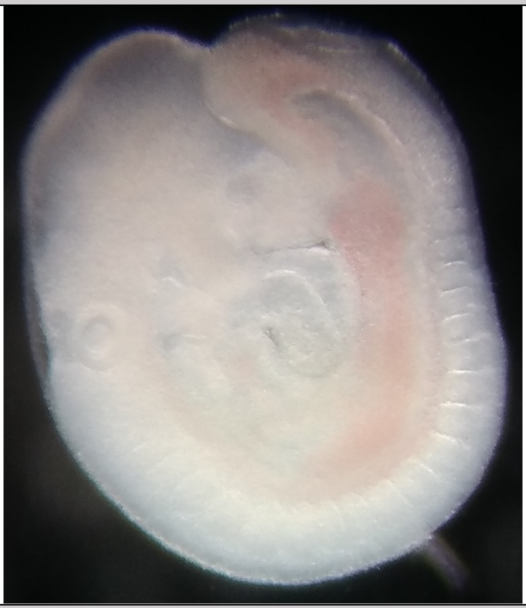 |
| **e15** | **e27** |
| 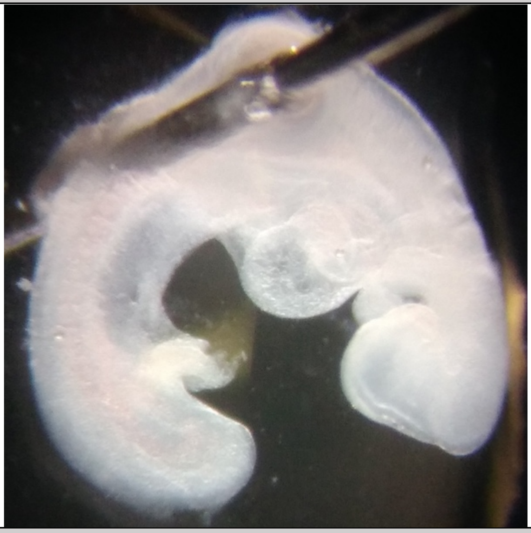 | 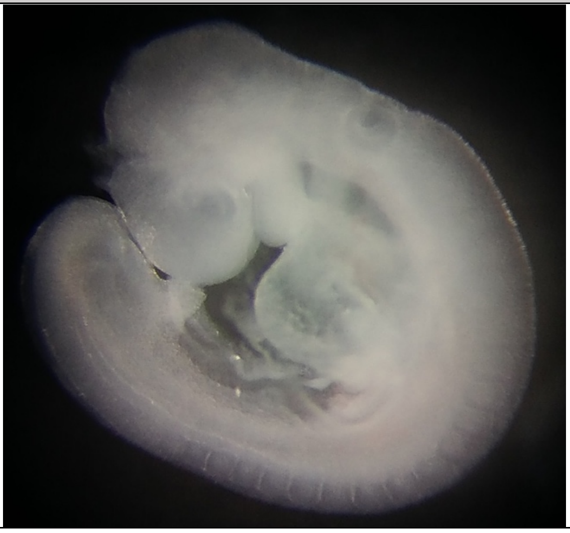 |
| **e28** | |
| 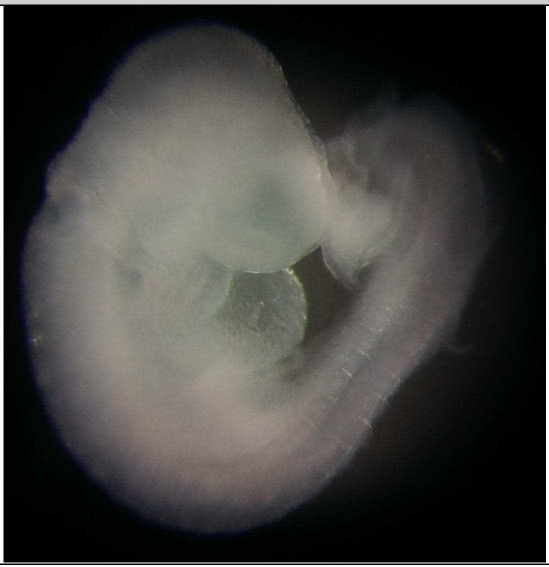 | |

### Stage: 21 somite pairs

| e1 | e2 |
|---|---|
| 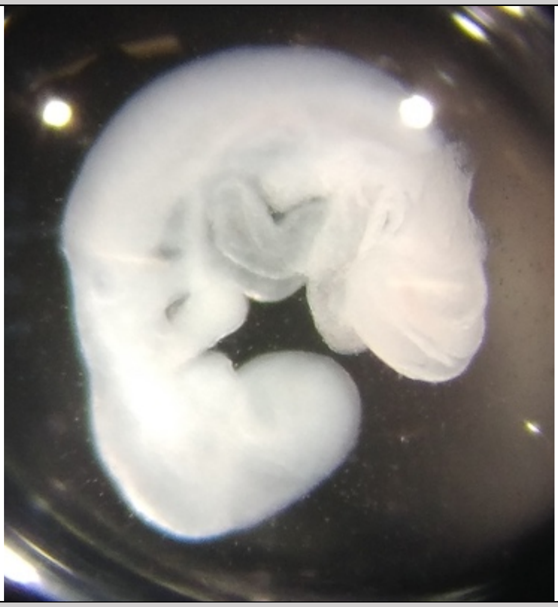 | 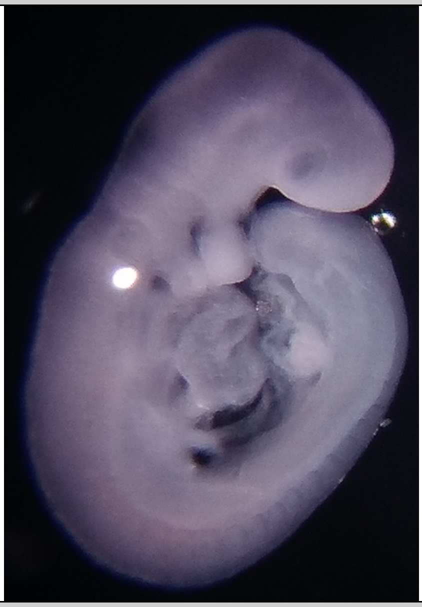 |
| **e3** | **e13** |
| 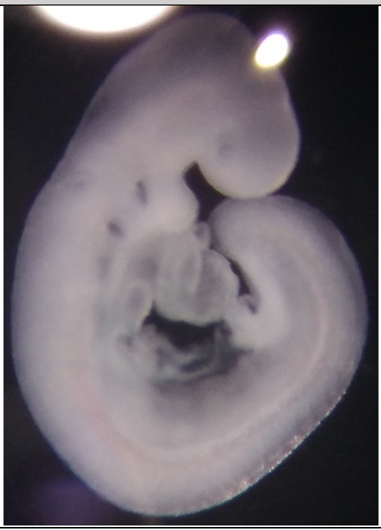 | 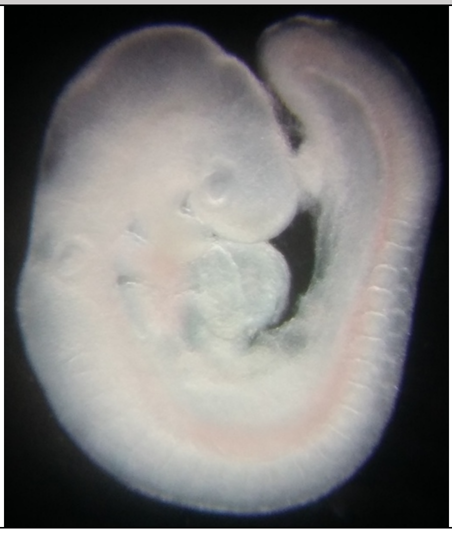 |

### Stage: 25 somite pairs

| e19 | e20 |
| --- | --- |
| 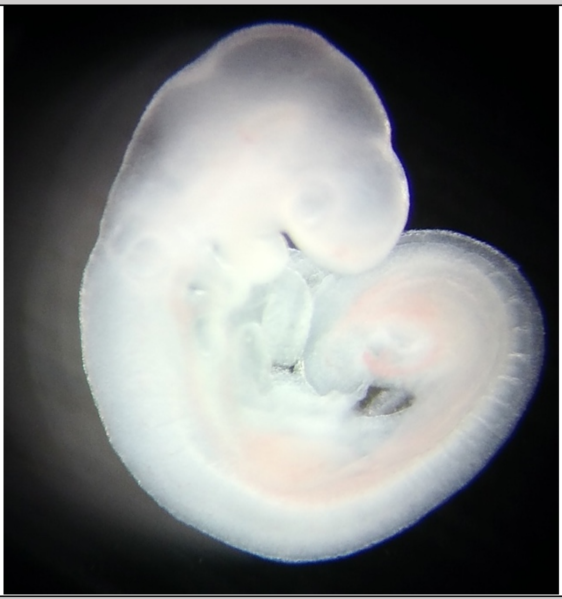 | 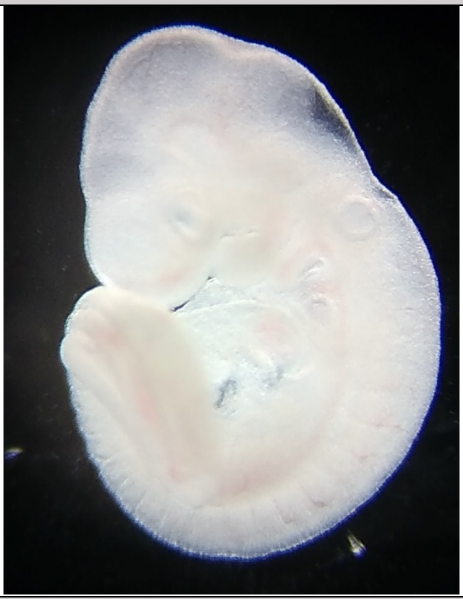 |
| **e21** | **e22** |
| 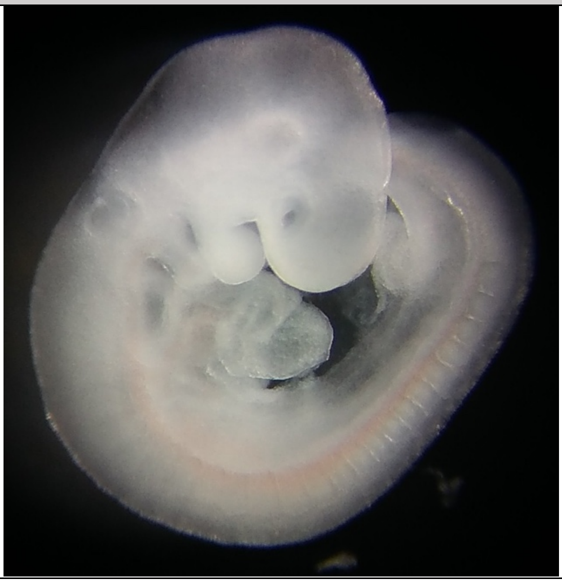 | 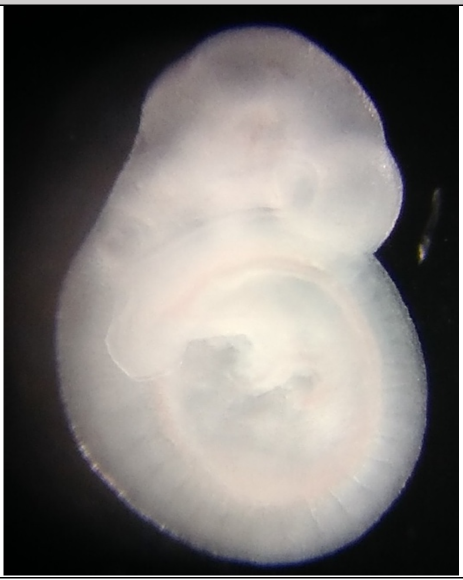 |

### Stage: 27 somite pairs

| e23 | e24 |
|---|---|
| 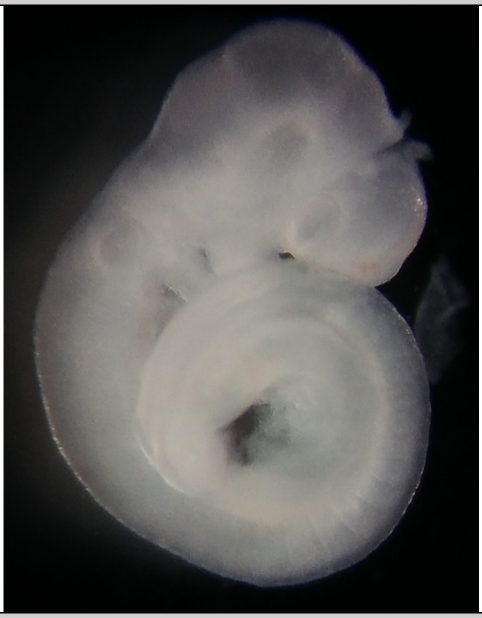 | 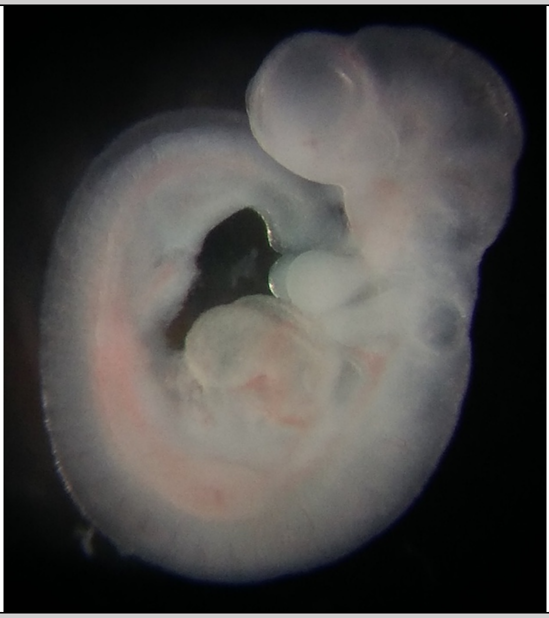 |
| **e25** | **e34** |
| 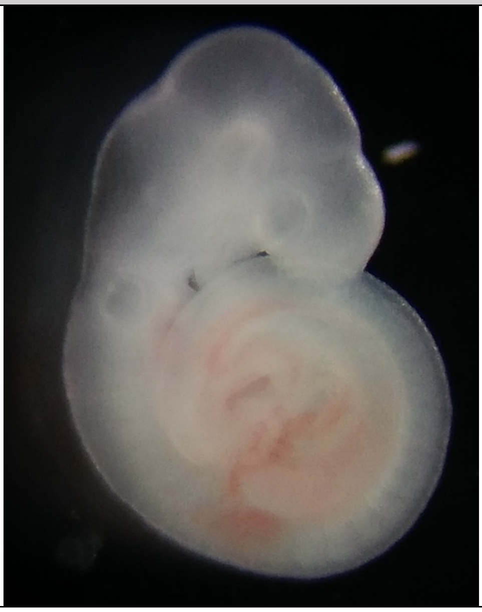 | 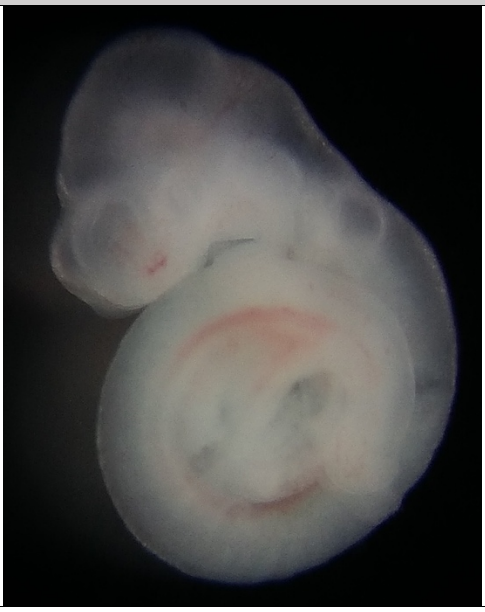 |

### Stage: 35 somite pairs

| e17 | e29 |
| --- | --- |
| 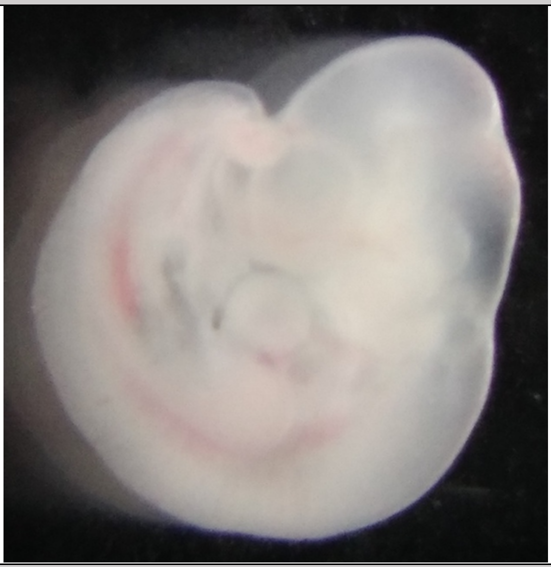 | 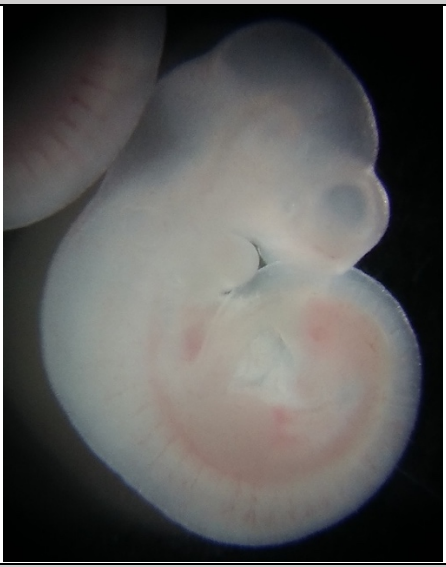 |
| **e30** | **e31** |
| 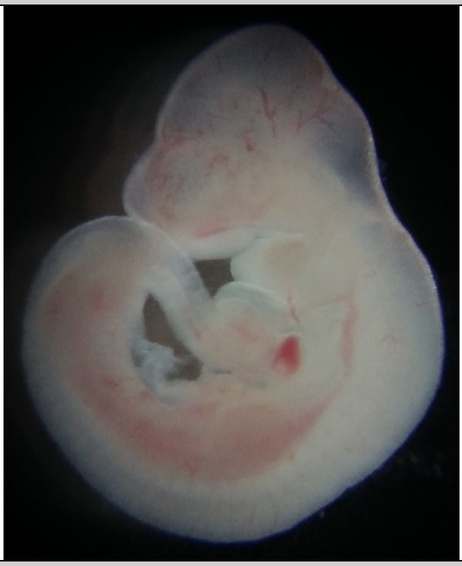 | 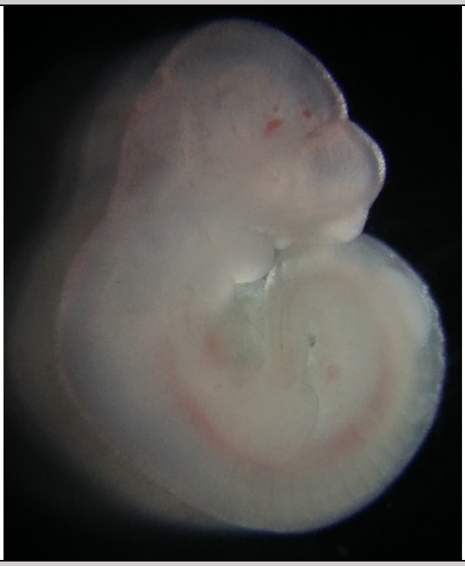 |
| **e32** | **e33** |
| 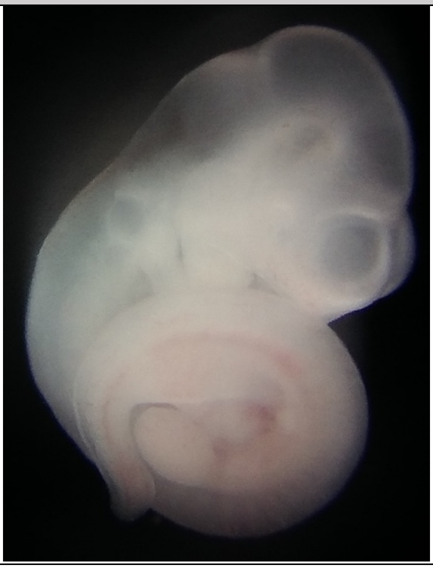 | 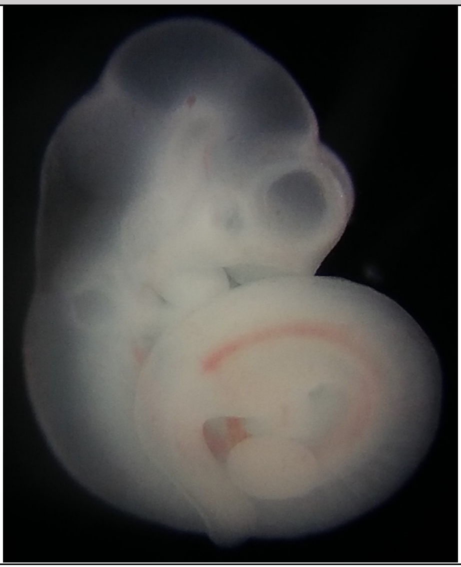 |
